# A role of oligodendrocytes in information processing independent of conduction velocity

**DOI:** 10.1101/736975

**Authors:** Sharlen Moore, Martin Meschkat, Torben Ruhwedel, Iva D. Tzvetanova, Andrea Trevisiol, Arne Battefeld, Kathrin Kusch, Maarten Kole, Nicola Strenzke, Wiebke Möbius, Livia de Hoz, Klaus-Armin Nave

## Abstract

Myelinating oligodendrocytes enable fast impulse propagation along axons as revealed through studies of homogeneously myelinated white matter tracts. However, gray matter myelination patterns are different, with sparsely myelinated sections leaving large portions of the axons naked. The consequences of this patchy myelination for oligodendrocyte function are not understood but suggest other roles in information processing beyond the regulation of axonal conduction velocity. Here, we analyzed the contribution of myelin to auditory information processing using paradigms that are good predictors of speech understanding in humans. We compared mice with different degrees of dysmyelination using acute cortical multiunit recordings in combination with behavioral readouts. We identified complex alterations of neuronal responses that reflect fatigue and temporal acuity deficits. Partially discriminable but overall similar deficits were observed in mice with oligodendrocytes that can myelinate but cannot fully support axons metabolically. Thus, myelination contributes to sustained stimulus perception in temporally complex paradigms, revealing a role of oligodendrocytes in the CNS beyond the increase of axonal conduction velocity.

## Introduction

In the central nervous system (CNS), oligodendrocytes generate myelin, a multilayered sheath of membrane, spirally wrapped around axonal segments and best known for its role in enabling fast saltatory impulse propagation ^1^. An additional function of oligodendrocytes is the metabolic support of myelinated axons ^2–4^, most important when axons spike at higher frequency ^5^. Mouse mutants with abnormalities of CNS myelination have been used as models of human leukodystrophies and demyelinating diseases. However, analyses of these mice have been largely restricted to the diagnosis and assessment of severely reduced motor functions.

Importantly, at the network level, the contribution of CNS myelination to information processing remains poorly understood, in part because suitable phenotyping instruments have been lacking. Yet, there is an increasing awareness that myelination is required for higher-order brain functions. Subtle defects of the white matter have been associated with various psychiatric diseases ^6^. Moreover, in the aging brain, the physiological decline of function is associated with subtle but widespread structural deterioration of myelin ^7,8^. Functional studies of myelin have largely been restricted to motor behavior, which is under control of myelinated axons in white matter tracts, and to the fast impulse propagation and millisecond precision required for sound location in a highly specialized circuit of the brain stem ^9^. However, myelin is also present within the cortical grey matter, where it is heterogeneous and patchy along projection neurons ^10,11^. Moreover, a large fraction of the parvalbumin-positive interneuron axons are myelinated ^12–14^, even though their typical short distance to their target cells has been estimated to cause insignificant conduction delay times ^13^. Oligodendrocyte lineage cells can myelinate axons in an activity dependent manner and influence their conduction velocity properties ^15–19^. Mature oligodendrocytes respond to glutamatergic signals with enhanced glycolytic support of the axonal energy metabolism ^4,5^. These data are compatible with the idea that myelin function in the CNS that go beyond regulating conduction velocity.

Indeed, there is still a major conceptual gap between the role of myelin in saltatory impulse propagation and its function at the network level. Virtually nothing is known about the contribution of myelination to information processing, a function that should be most evident for pathways that build on constant information transfer, as it is the case for the auditory system. In the sound location system, nerve conduction velocity is fine-tuned by myelination beginning at the brainstem level in order to facilitate sound localization ^20,21^. However, the sound location system is a highly specialized circuit dedicated to sub-millisecond precision of coincidence detection. Auditory processing is clearly broader than sound localization and must constantly cope with the appearance of both short and repetitive stimuli. This is essential e.g. for speech perception.

In humans, myelin abnormalities and nerve conduction velocities have been mostly studied in terms of developmental delays, and rarely with respect to signal detection, precision and acuity. And yet, it is clinically well known that patients with myelin disease, similar to individuals at old age, show deficits in auditory processing ^22,23^ that cannot simply be explained by delays in conduction velocity. For example, deficits in speech recognition in noisy environments increase despite normal hearing thresholds ^24^, which is suggestive of problems in temporal acuity, rather than the speed of processing.

In animal models of CNS dysmyelination, a few studies reported general auditory abnormalities ^25–28^ or a specific delay at central auditory stations ^29–31^, as detected by evoked response latencies. Using brainstem slices, e.g. Kim et al. reported excitability defects at the Calyx of Held, a specialized synapse in the brainstem, important for sound localization ^28^. However, the perceptual consequences have not and cannot be assessed with an *in vitro* system.

Here, we used a novel approach for the *in vivo* analysis of information processing and perception in mice as a function of oligodendrocyte and myelin abnormalities. To determine how dysmyelination and impairments in axonal metabolic support affect temporal and spectral auditory processing, we compared adult wild type animals with three lines of mutant mice. We used 1) the *shiverer* mice (*Mbp*^*shi/shi*^) ^32^ as a model of severe dysmyelination, 2) a newly generated *Mbp* allele (*Mbp*^*neo/neo*^) as a hypomyelination model (see below), and 3) mice with the heterozygous null allele of the *Slc16A1* gene (MCT1^*+/-*^) ^3^, encoding a monocarboxylate transporter (MCT1) that is required for metabolic support. The use of these mouse models allowed the comparison of different degrees of myelination, and to distinguish that from the role of oligodendrocytes in axonal metabolic support.

Multiunit recordings of the cortex enabled us to test stimulus detection during auditory processing in combination with behavioral readouts. Altered neuronal responses in the auditory cortex of dysmyelinated mice were surprisingly complex, and included neuronal fatigue, likely energy-related, and temporal acuity deficits that clearly exceeded the effect of delays in conduction velocity. Moreover, deficits in “sound gap detection”, a measure relevant to speech recognition in humans, indicate that myelination of CNS axons is critical for normal sound processing beyond the (data not shown) finely tuned conduction velocity for sound localization. Our data reveal that CNS myelination not only speeds impulse conduction but also determines the quality of information processing, which we suggest is partially dependent on the metabolic roles of oligodendrocytes.

## Results

To monitor auditory processing in the adult brain with compromised oligodendrocyte functions, we chose three types of mutant mice. To study the role of compacted myelin, we compared different *Mbp* mutations in mice. MBP is an abundant structural protein required for myelin growth and compaction ^33,34^. One mutant is defined by the nearly complete absence of CNS myelin, the well-known *shiverer* (*Mbp*^*shi/shi*^) mouse ^35^ with a truncated *Mbp* gene ^32^. The second *Mbp* model was newly generated for this study as a hypomorph mouse (*Mbp*^*neo/neo*^) with MBP expression levels below 50% as detailed below. A third mutant, heterozygous null for the *Slc16A1* gene (MCT1^*+/-*^) shows reduced expression of the monocarboxylate transporter MCT1, which is required by oligodendrocytes to metabolically support axons ^3^. MCT1 enables the export of lactate and pyruvate and supports the generation of ATP in myelinated axons ^5^.

### *Mbp^neo^*: a novel mouse mutant with reduced myelin sheath thickness

Shiverer mice have no compact myelin and a shivering phenotype that makes them suboptimal for behavioral testing. Since heterozygous *Mbp*^*shi/+*^ mice show close to normal amounts of myelin ^36^, we sought to establish a hypomyelinated mouse model by reducing *Mbp* expression levels below 50% (see methods and Figure S1A), in agreement with earlier observations of transgenic complementation of *shiverer* mice ^37,38^. Oligodendrocytes in adult *Mbp*^*neo/neo*^ brains expressed 30% of the *Mbp* mRNA and 20% MBP at the protein level (Figures S1B and S1C). These mutants appeared clinically normal and were long-lived. Different from conventional *shivere*r mice, in which the vast majority of optic nerve axons are unmyelinated ^35^, *Mbp*^*neo/neo*^ mice have 75% of their optic nerve axons myelinated (wild type: 92%). Importantly, this myelin was fully compacted but significantly thinner, as determined by g-ratio analysis (Figures S1D to S1F). Axon caliber distribution was not different between wild type and *Mbp*^*neo/neo*^ mice (Figure S1G).

### Central dysmyelination causes signs of auditory neuropathy

In wild type mice, all auditory projections, including the auditory nerve, are well myelinated ^39,40^. Before studying the effect of dysmyelination on information processing, we determined latencies and amplitudes of sound-evoked responses at different auditory stations using auditory brainstem responses (ABR), recorded *in vivo* through electrodes located on the scalp (Figure 1A i). ABRs consisted of five waves (I-V) that reflect the consecutive activation of the auditory nerve, cochlear nucleus, superior olive, lateral lemniscus and inferior colliculus (Figure 1A ii). In the *Mbp*^*shi/shi*^ mice, ABR waves were significantly delayed at all auditory stations (Figures 1B and 1C) and delays increased progressively. The peak-to-peak amplitude, which reflects the synchronous spiking in response to sound onset, was unchanged in wave I (Figure 1F). This, together with the findings that also response thresholds were not different from controls (Figures S2A and S2B), confirms that cochlear function and peripheral conduction is normal in *Mbp*^*shi/shi*^ mice. This was expected because the outer region of the auditory nerve is myelinated by Schwann cells that do not require MBP for myelination (Figure 1A iii) ^40^. However, the amplitudes of waves II and III were reduced, and those of waves IV and V were increased (Figures 1B and 1F). Such downstream improvements of diminished brainstem ABR amplitudes have been observed in several animal models of auditory synaptopathy/neuropathy (disorders of the inner hair cell ribbon synapse and/or auditory nerve), and are thought to reflect compensatory mechanism of changes in gain to peripheral hearing loss ^41–46^. Hypomyelinated *Mbp*^*neo/neo*^ mice were similarly normal with respect to the shape of ABR wave I (Figures 1D and 1E) and threshold (Figure S2C). They also showed a reduction in the amplitudes of waves II and III, and a larger wave V at the IC level (Figure 1G). The *Mbp*^*shi/shi*^ and *Mbp*^*neo/neo*^ wave V/I ratio increase appeared proportional to the degree of dysmyelination (230% and 100% increase respectively; Figure 1F and 1G, bottom right). In contrast to the dysmyelination models, MCT1^*+/-*^ mutants revealed no changes in threshold, latency, or amplitude of ABR waves (Figures S2E to S2H). Thus, latency increase, and signal desynchronization are a result of partial or complete dysmyelination, unrelated to metabolic defects.

**Figure 1.**
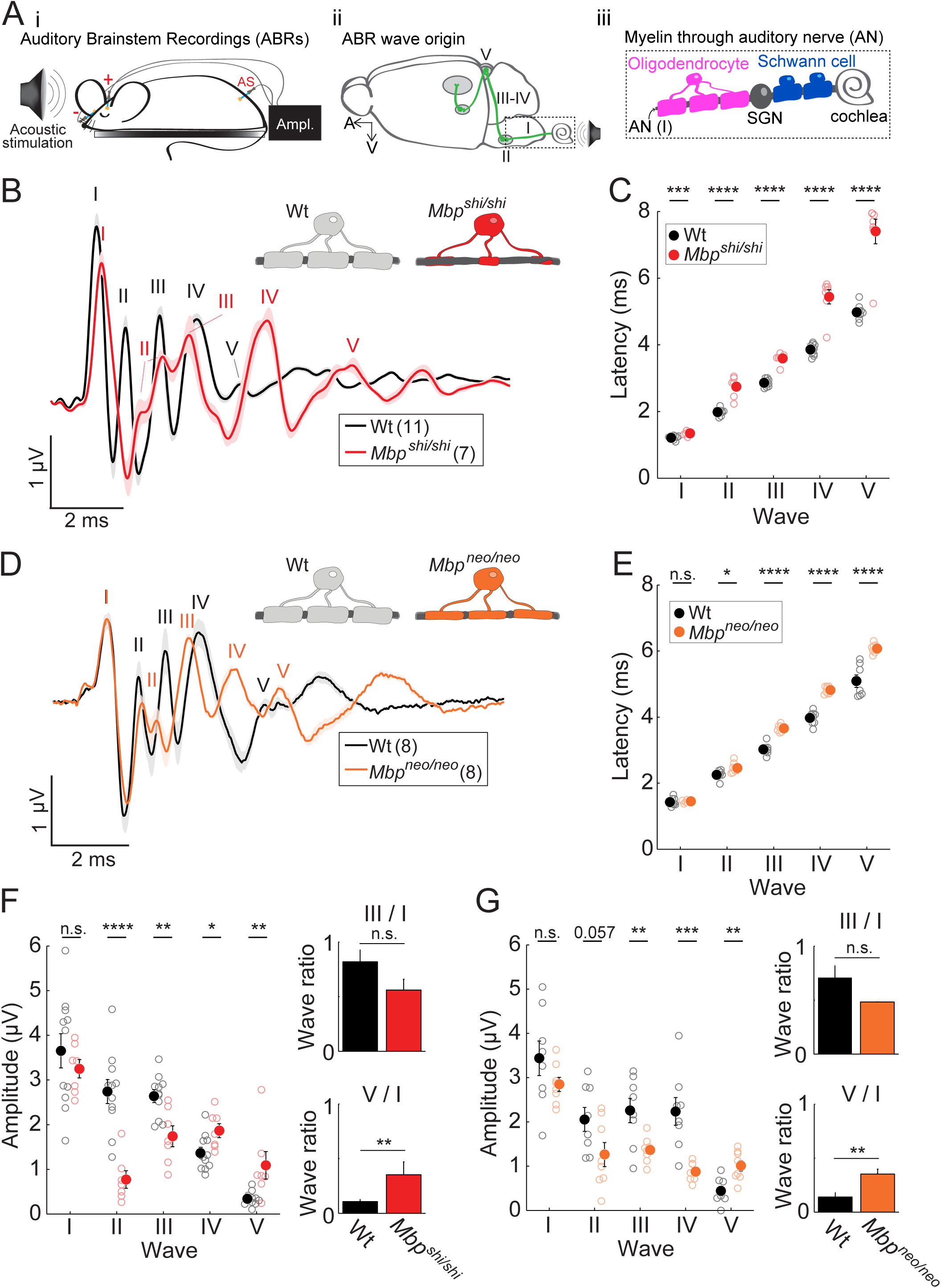
Central dysmyelination causes evoked-response delays and gain changes. **A**) Left (i): Schematic of auditory brainstem (ABR) scalp recordings in mice. Middle (ii): structures in the auditory circuit contributing to ABRs (I to V). Right (iii): scheme of peripheral (around the spiral ganglion neurons, SGN) and central myelination of auditory nerve (AN). **B)** Group mean traces in control (black; n=11; 7 *Mbp*^+/+^ and 4 *Mbp*^*shi/+*^ without significant differences in ABR measurements (80 dB), see table in Figure S2D) and *Mbp*^*shi/shi*^ (red; n=7) mice. Each one of the 5 peaks (I–V) can be attributed to activity at a different station along the auditory brainstem (see schematic in Aii). Responses in *Mbp*^*shi/shi*^ mice were delayed at all auditory stations. Wave II appears divided and merged with wave III. **C)** The quantification of the peak latencies reveals a progressively larger difference in latency from wave I (p=0.001) to wave V (p<0.0001 for waves II-V). **D)** Same as A) but for *Mbp*^*neo/neo*^ mice (orange; n = 8) and Wt (black; n=8). Wave II appears divided. **E)** Latency quantification again showed progressively larger latency differences from wave I to wave V (p=0.74, p=0.013, p<0.0001, p<0.0001, p=0.0001; waves I-V, respectively). **F)** Wave amplitude quantification. In *Mbp*^*shi/shi*^ mice (red), waves II and III amplitudes were reduced, while those of waves IV and V were increased (p=0.44, p<0.0001, p=0.0031, p=0.022, p=0.008; waves I to V respectively), suggesting a compensatory central gain increase. Auditory gain was quantified by the amplitude ratio in waves III/I (upper right graph), which was comparable across groups (p=0.085), and waves V/I (lower right graph), which was significantly increased in mutants (p=0.006). **G)** Same as E but for *Mbp*^*neo/neo*^ mice. As in *Mbp*^*shi/shi*^ mice, amplitude changes were seen at the central level (p=0.18, p=0.057, p=0.009, p=0.0008, p=0.003; waves I to V respectively). An increase in central gain was observed in *Mbp*^*neo/neo*^ for wave V (lower right graph; p=0.009) but no changes in the gain of wave III (upper right graph; p=0.083). All graphs depict the mean and S.E.M. (error bar or a shaded area).

### Myelination of primary auditory cortex is patchy and heterogeneous

To study auditory processing, we first assessed the presence of myelin in the inferior colliculus (IC), an important subcortical relay station in the auditory path, and in layer 4 of the primary auditory cortex (ACx), where we obtained multi-unit recordings (Figure 2A). As in other grey matter areas, myelination profiles are present at low density ^10,13^. Compared to the IC, myelination of the ACx was even sparser (Figure 2B, left upper panel). In *Mbp*^*shi/shi*^ mice there was no compact myelin, but oligodendrocyte processes loosely ensheathed the axons. Interestingly, when quantified, both *Mbp*^*shi/shi*^ (*shiverer*) and *Mbp*^*neo/neo*^ mice had the same number of ensheathed axons (quantified irrespective of their compaction) in the ACx as their respective controls (Figure 2B, right panels graphs). Dysmyelination of *Mbp*^*neo/neo*^ was only obvious by electron microscopy (Figure 2B, lower insets). Wild type axons were surrounded by electron-dense (compact) myelin (Figure 2B, left panels), *Mbp*^*shi/shi*^ axons were loosely ensheathed (Figure 2B, right upper panel) and *Mbp*^*neo/neo*^ mice exhibited an intermediate profile with compact but thinner myelin compared to wild type (Figure 2B, right lower panel).

**Figure 2.**
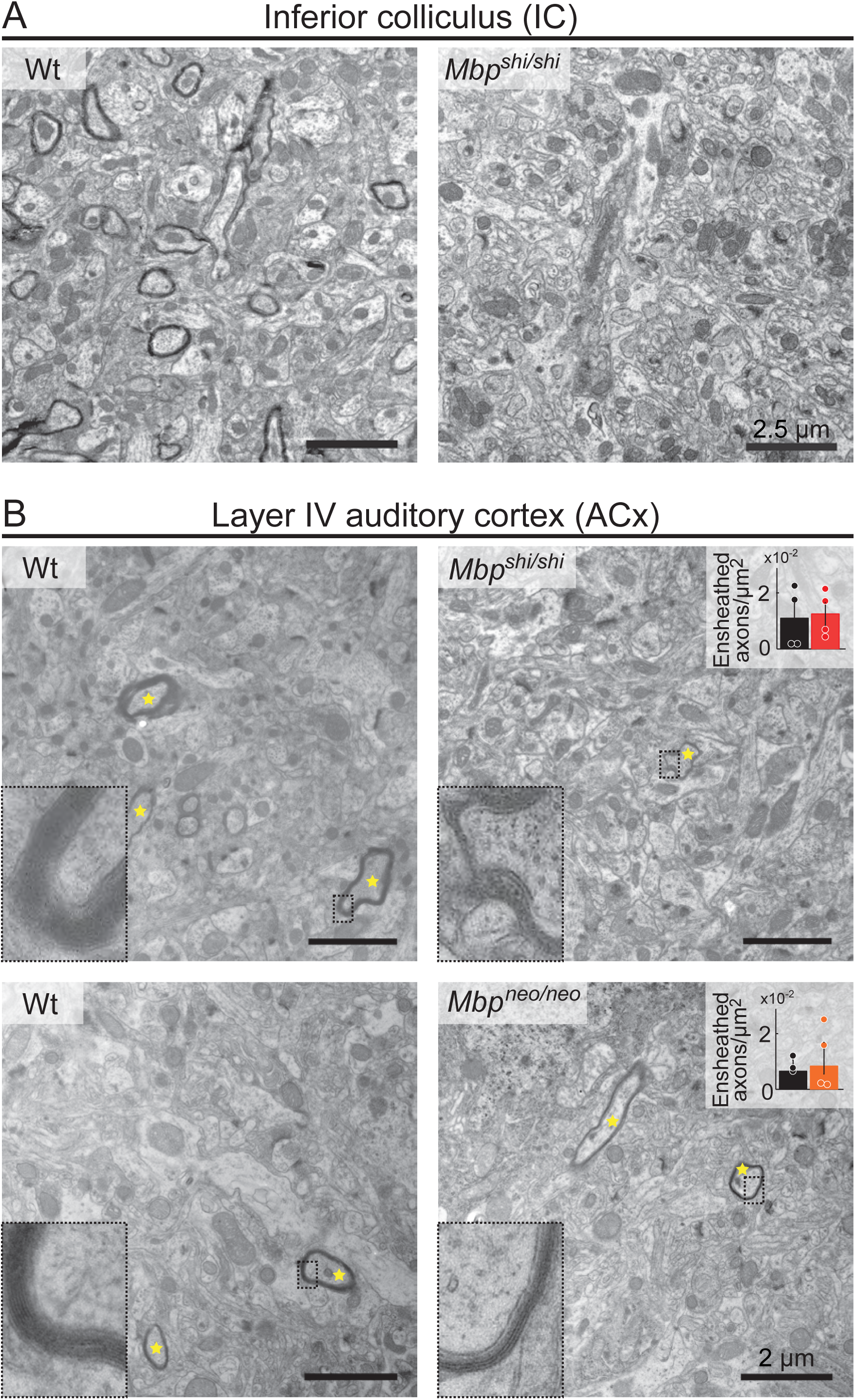
Ensheathment profiles along the auditory system in normal and dysmyelinated models. **A)** Electron microscopy images of the inferior colliculus of a Wt mouse (left panel) showing sparse compact myelin, and an *Mbp*^*shi/shi*^ mouse (right panel), lacking electro-dense compact myelin. **B)** Electron microscopy images of the auditory cortex of Wt and *Mbp*^*shi/shi*^ (upper panel) and Wt and *Mbp*^*neo/neo*^ mouse (lower panel). Ensheathed axons in the ACx are marked with yellow asterisks. Insets show details of the myelin sheath of axons from the respective image. *Mbp*^*shi/shi*^ axons (upper right) show lack of compact myelin, while *Mbp*^*neo/neo*^ axons (lower right) show thinner compact myelin than Wts. Inset plots show the quantification of the number of ensheathed axons (compact or not compact myelin) per area in *Mbp*^*shi/shi*^ (left, red, n=4), *Mbp*^*neo/neo*^ (right, orange, n=4), and their respective Wts (black, n=3-4). There was no difference between Wt and either mutant model (p=0.83 and p=0.73 for *Mbp*^*shi/shi*^ and *Mbp*^*neo/neo*^ respectively). Bar graphs show the mean of all animals quantified (10-15 images per mouse). Scale bars: 2.5 µm (A), 2 µm (B).

### Cortical responses to simple sounds in dysmyelinated mice

To study the effect of dysmyelination, we performed *in vivo* extracellular recordings in *Mbp*^*shi/shi*^ mice (Figure 3A) following basic single sound stimuli (clicks) to evoke responses. Surprisingly, at the level of the IC, we obtained an increased response strength compared to controls (Figures 3B and S3A). However, at the level of the ACx, response strength was similar to controls both in *Mbp*^*shi/shi*^ (Figures 3C and S3B, left) and *Mbp*^*neo/neo*^ mice (Figures 3D and S3B, middle). In contrast, response latencies (Figure 3E) were increased in *Mbp*^*shi/shi*^ at both auditory stations (Figures 3C, 3F and 3G) similar to *Mbp*^*neo/neo*^ mice (Figures 3D and 3H), and this increase correlated with the degree of dysmyelination (Figure S3B right). Response jitter (Figure S3C) and inter-trial reliability (Figure S3G) were not different in any of the models or stations (Figure S3D to S3F and S3H to S3J). Not surprisingly, cortical neurons in the normally myelinated MCT1^+/-^ mice responded like controls to simple clicks and showed only a tendency towards longer latencies (Figure S3K to S3N).

**Figure 3.**
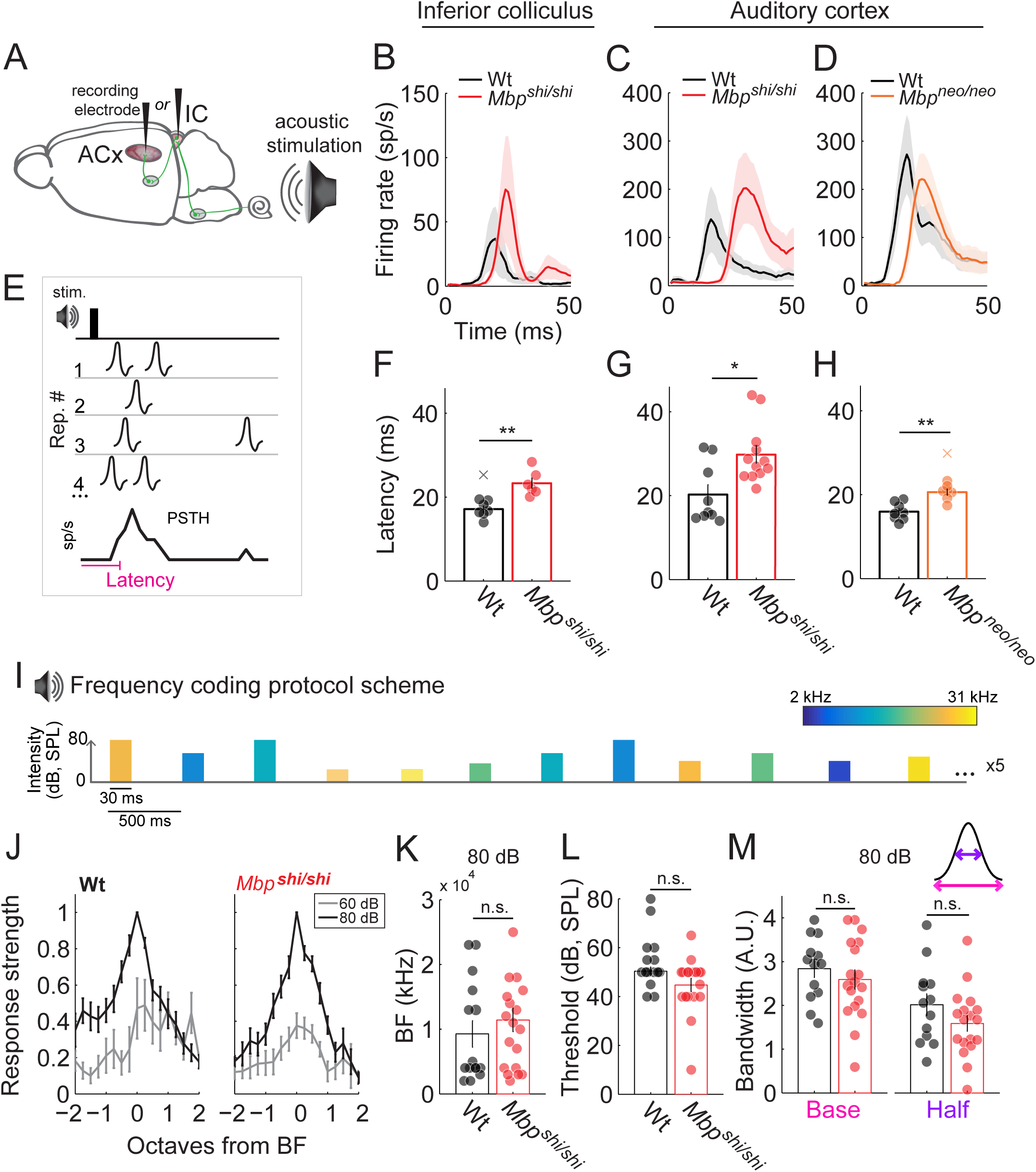
Responses to single sound-stimuli in IC and ACx of dysmyelinated models are abnormal only in latency. **A)** Schematic brain depicting auditory areas (ACx and IC) where multi-unit recordings took place. **B)** Peri-stimulus time histogram (PSTH) of responses to a click sound recorded in the IC for Wt (black; n=6-8) and *Mbp*^*shi/shi*^ (red; n=6-8) mice aligned at time of click presentation (0 ms). Responses of *Mbp*^*shi/shi*^ are delayed with respect to Wt. **C)** Same as B) but for ACx recordings. A stronger delay in *Mbp*^*shi/shi*^ (red; n=12-13) compared to Wt (black; n=7-10) is observed in this case. **D)** Same as C) but for *Mbp*^*neo/neo*^ mice (orange; n=7-8) and corresponding Wt (black; n=8). A smaller shift in latency in the ACx of *Mbp*^*neo/neo*^ is observed, compared to *Mbp*^*shi/shi*^ mice. **E)** Scheme of latency measurement: time at which the PSTH, sum of spikes across 10 trials in 1 ms time windows, surpassed 1.5 times the baseline activity. A significant latency increase is seen in *Mbp*^*shi/shi*^ animals compared to Wt. **F)** in the IC (p=0.0012) and **G)** in the ACx (p=0.023). **H)** *Mbp*^*neo/neo*^ mice also showed a significant increase in response latency in the ACx (p=0.0012). **I)** Schematic of the tone sweep protocol used to test tuning. A series of 24, 30 ms long, tones (2-31 kHz) were played at different intensities. 10 repetitions (Rep.) of each tone-intensity combination were presented in a random order. **J-M**) Basic tuning properties are not affected with dysmyelination. n=13 recordings of 10 wild type mice. n=18 recordings of 15 *Mbp*^*shi/shi*^ mice. **J)** Normalized tuning curves for wild type (left) and *Mbp*^*shi/shi*^ mice show response selectivity in octaves from best frequency (BF) at 60 (gray) and 80 dB (black). **K)** Recordings from comparable rostro-caudal locations yielded comparable BFs at 80 dB (p=0.51) in wild type and *Mbp*^*shi/shi*^ mice indicating no major change in tonotopic map. **L)** Auditory cortical thresholds were similar between groups (p=0.11). **M)** Bandwidth of tuning was also similar between groups (p=0.56 and p=0.21 for base and half bandwidth respectively).

Changes in the ion channel distribution of the axon initial segment (AIS) are believed to underlie dysmyelination-dependent changes of excitability ^47^. Puzzled by the fact that the hyperexcitable responses detected in the IC of *Mbp*^*shi/shi*^ mice were not present in the ACx (Figure 3B to 3C and S3A to S3B), we measured AIS length in the ACx. Interestingly, here *Mbp*^*shi/shi*^ mice exhibited a reduction in the lengths of AIS (Figures S4A and S4B), which may explain the absence of hyperexcitable responses, as a short AIS is typically associated with reduced intrinsic neuronal excitability ^47–49^.

### Spectral processing of pure tones

Key features of spectral processing are frequency tuning and response adaptation, both of which are strongly influenced by neuronal integration, i.e. the functional convergence of stimuli onto individual cortical neurons. Frequency tuning, a property of the auditory system, builds on frequency-specific inputs onto a given cell and is modulated by lateral inhibition. Thus, differential conduction velocities in converging pathways as a result of reduced myelination could have a lasting effect on neuronal integration. Surprisingly, upon stimulation with pure tones (Figure 3I), tuning curves had comparable shapes in *Mbp*^*shi/shi*^ and wild type mice, as a function of sound intensity (Figure 3J), and covered comparable best-frequency (BF) ranges within the sampled regions (Figure 3K). There was no difference between groups in hearing thresholds (Figure 3L) or tuning bandwidth (Figure 3M), which depends largely on convergent inputs ^50^.

Adaptation is crucial for sensory filtering in the auditory system ^51,52^ and a measure of neuronal integration. It is typically tested using oddball paradigms (Figure S4C), in which two frequencies that elicit responses of similar magnitude are presented sequentially such that one appears with higher probability (‘standard’) than the other (‘deviant’). Here, stimulus-specific adaptation (SSA) is reflected in a decreased response to the standard tone, while the response to the deviant tone remains constant or increases ^52,53^. Importantly, cortical SSA was not reduced in *Mbp*^*shi/shi*^ compared to wild type mice (Figure S4D). Thus, widespread dysmyelination had no obvious effect on the integration of auditory stimuli, presumably because the convergent inputs were all similarly delayed.

### Dysmyelination affects temporal reliability and acuity of sound processing

Naturally occurring sounds are characterized by rich temporal structures. In many species, the reliable temporal coding of communication sounds is an important factor for survival or successful mating. Correct auditory coding relies on the ability of the nervous system to convey the temporal structure of sounds with precise spike timing and to maintain this precision while listening to continuous sound streams. We determined, on one hand the neuronal capacity of our mouse models to follow click-trains (a measure of temporal reliability) at different presentation rates. We also determined ‘gap-detection’, a measure of temporal acuity, as used for audiometric testing in humans ^54–56^.

Click-rate detection was tested with a set of 10 clicks presented at different temporal rates (Figure 4A). In wild type mice, the rates at which auditory neurons can follow were lower in ACx compared to subcortical stations, consistent with a previous report ^57^. Repeated presentations revealed that the timing of spikes is reproducible, although failures were observed in the response to the last clicks in this sequence (Figure 4B, top). Importantly, in *Mbp*^*shi/shi*^ mice, ACx neurons followed the initial clicks in a set of 10, but (already in the 5 Hz condition) revealed a strong decay in the response strength to the last clicks (individual example Figure 4B, bottom). Scoring the responses to click number 1, 5 and 10 (taken as representatives), we observed that the response magnitude in *Mbp*^*shi/shi*^ mice was the same as in wild type after click 1 but was smaller after click 5 and virtually non-existent by click 10 (Figure 4C). This decrease in response magnitude was associated with a decreased synchrony of spiking (between repetitions along the 10 clicks) compared to wild type (Figure 4D and 4E). A similar effect was observed in the cortex of *Mbp*^*neo/neo*^ mice (Figure 4E center). Thus, ‘naked’ axons of *Mbp*^*shi/shi*^ mice were not a prerequisite for this observation. Importantly, MCT1^*+/-*^ mice, which are normally myelinated but fail to fully support axonal energy metabolism, exhibited the same deficit, which was evident at even slower click rates (Figure 4E right). This suggests that axonal energy deficits are the likely cause of conduction failure following repetitive neuronal firing.

**Figure 4.**
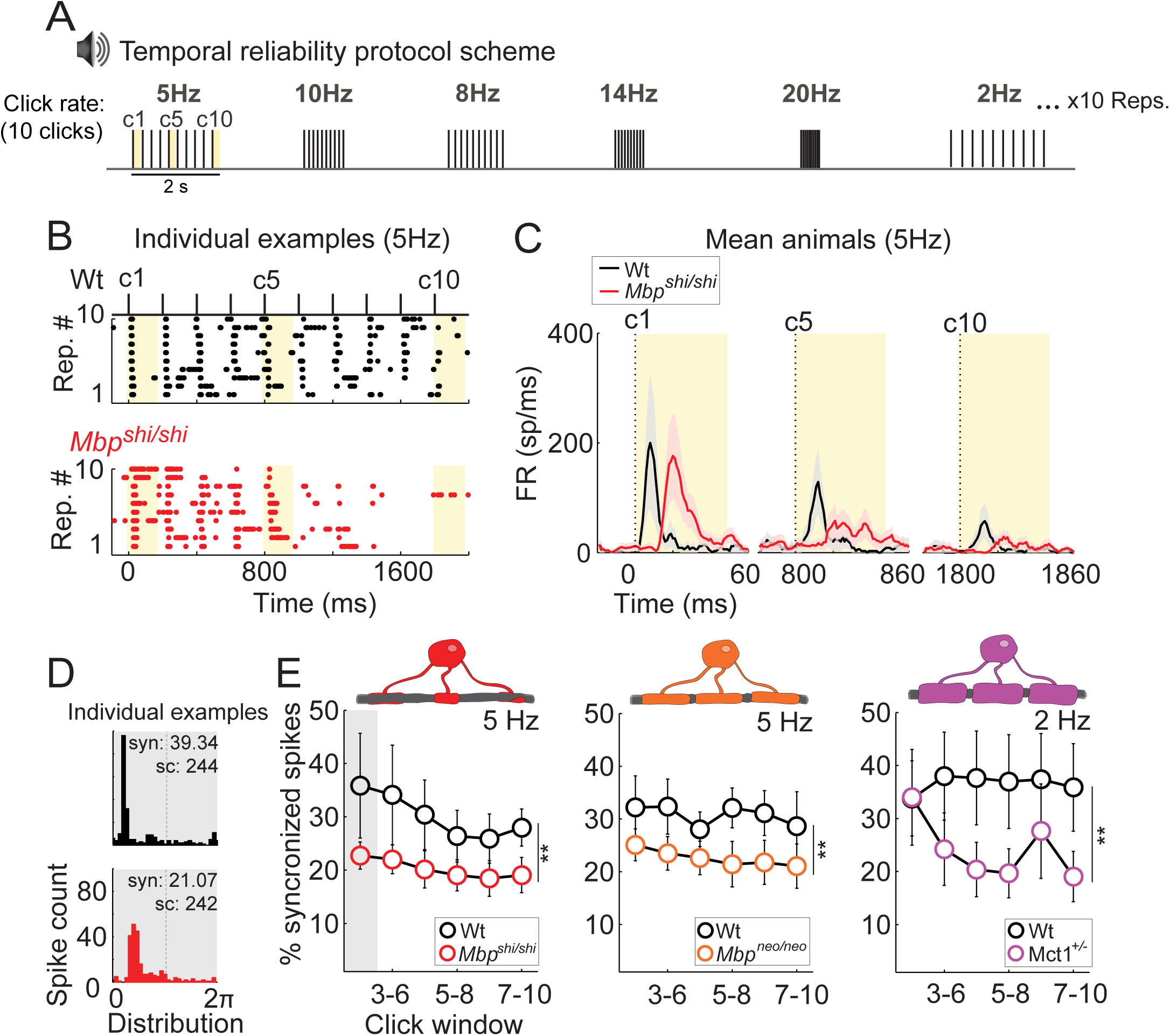
Temporal reliability is affected in mice with either dysmyelination or a reduction in axo-glial metabolic support. **A)** Schematic of the click rate-tracing protocol used to test temporal reliability. Blocks of 10 clicks are played at different rates in random order. Each rate is repeated 10 times. Analysis in C) focuses on click 1, 5 and 10 (c1, c5 and c10). **B)** Example raster plot of ACx responses (each dot represents 1 spike) to 10 clicks at 5 Hz across the 10 stimulus repetitions in a Wt (upper, black) and an *Mbp*^*shi/shi*^ mouse (lower, red). *Mbp*^*shi/shi*^ animals show a steeper decay on spiking activity across clicks compared to Wt. **C)** Mean PSTH of responses to click 1, 5 and 10, at 5 Hz for Wt (black; n=10) and *Mbp*^*shi/shi*^ (red; n=13) animals. The thick line shows the mean of all recorded animals and the shaded area depicts the S.E.M. Click time is indicated by dashed lines. While responses to the first click are similar in amplitude in Wt and *Mbp*^*shi/shi*^ animals (with the expected delay in *Mbp*^*shi/shi*^), a strong reduction of response strength is seen in *Mbp*^*shi/shi*^ mice with increasing clicks. **D)** Individual examples of spike synchrony plots for Wt (top) and *Mbp*^*shi/shi*^ (bottom) were taken from the first click window (clicks 2-5, depicted by a gray square in E). **E)** Quantification of spike synchrony in sliding windows of 4 clicks (clicks 2-5, 3-6, 4-7, 5-8, 6-9 and 7-10). Onset responses of click 1 were not used. Leftmost: significant reduction in spike synchrony between *Mbp*^*shi/shi*^ mice (red, n=6-9) and Wt (black, n= 3-6 mice) at 5 Hz (p=0.0021). Middle: significant reduction also seen between *Mbp*^*neo/neo*^ mice (orange, n=5-8 mice), and Wt (black, n=3-8 mice) at 5 Hz (p=0.0014). Right: significant difference between MCT1^*+/-*^ mice (purple, n=5-6 mice), and Wt (black, n=8 mice) at 2 Hz (p=0.011).

The observed deficit in temporal reliability could be explained by the altered K_v_ channels distribution (dispersed along internodes and overexpressed in *Mbp*^*shi/shi*^) in axonal excitable domains ^58^, which we confirmed in the form of shortened AIS for the ACx of *Mbp*^*shi/shi*^ mice at the level of the AIS (Figures S4A and S4B). In a dysmyelinated brain, axonal excitability, membrane repolarization, and energy consumption could be affected by the general misdistribution of ion channels ^47,59^. As a model for other CNS white matter tracts that are difficult to study directly, we used a well-established optic nerve preparation ^60^. This acute *ex vivo* system allowed us to measure compound action potentials (CAP) for myelinated axons (Figure S5A) and to apply pharmacological manipulations ^4,5,60^. In wild type optic nerves (ON), CAPs exhibited the expected 3-peak shape, but we observed strong differences with both *Mbp*^*shi/shi*^ and *Mbp*^*neo/neo*^ nerves. In these mutants, CAP peaks were delayed in latency (Figure S5B) reflecting slowed nerve conduction velocity (Figure S5C) proportional to the level of dysmyelination (Figures S5D). Dysmyelinated optic nerves also showed a decrease in peak amplitude (Figure S5B) and CAP area (Figure S5G), suggesting conduction blocks in the ON, and we noted an increase of the hyperpolarizing phase with a larger negative CAP area (Figure S5H). These features were similar to CAP recordings from the spinal cord of *Mbp*^*shi/shi*^ mice ^58^. To test our hypothesis that excess potassium fluxes in dysmyelinated *Mbp*^*shi/shi*^ optic nerve axons cause energy loss (due to repolarization-associated ATP consumption) and contribute to reduced temporal reliability, we blocked potassium channels with 4-aminopyridine (4-AP, 25 µM) in a bath application. This treatment normalized the CAP amplitude and decreased the hyperpolarizing phase of *Mbp*^*shi/shi*^ axons (Figures S5F gray, S5G rightmost and S5H rightmost). This finding is compatible with abnormal ion fluxes in the mutant axons, which increase ATP consumption and cause abnormal repolarization dynamics.

The second key aspect of auditory processing is temporal acuity. Gap-detection protocols measure the ability of neurons to detect short silent gaps in the middle of a white noise sound (Figure 5A), a capacity also crucial in human speech processing ^61^. In mice, the presence and the strength of any post-gap sound response reflects whether the gap has been detected (Figure 5B). *Mbp*^*shi/shi*^ mice revealed a significant decrease (compared to wild types) in their post-gap response strength for gaps shorter than 3 ms in duration (Figures 5B and 5D). Quantification of this effect (by comparing for each recording the ratio between baseline activity and post-gap activity) confirmed that the cortical neurons of *Mbp*^*shi/shi*^ mice failed to detect small gaps (Figure 5C). A milder but similar effect was observed in *Mbp*^*neo/neo*^ mice (Figures 5E, S6A and S6B), suggesting that these temporal acuity deficits reflect the level of dysmyelination. Importantly, MCT1^+/-^ mice revealed only very minor gap detection deficit compared to their respective controls (Figures S6C to S6E). Thus, loss of temporal acuity must be attributed to hypomyelination, rather than reduced metabolic support of the axonal compartment.

**Figure 5.**
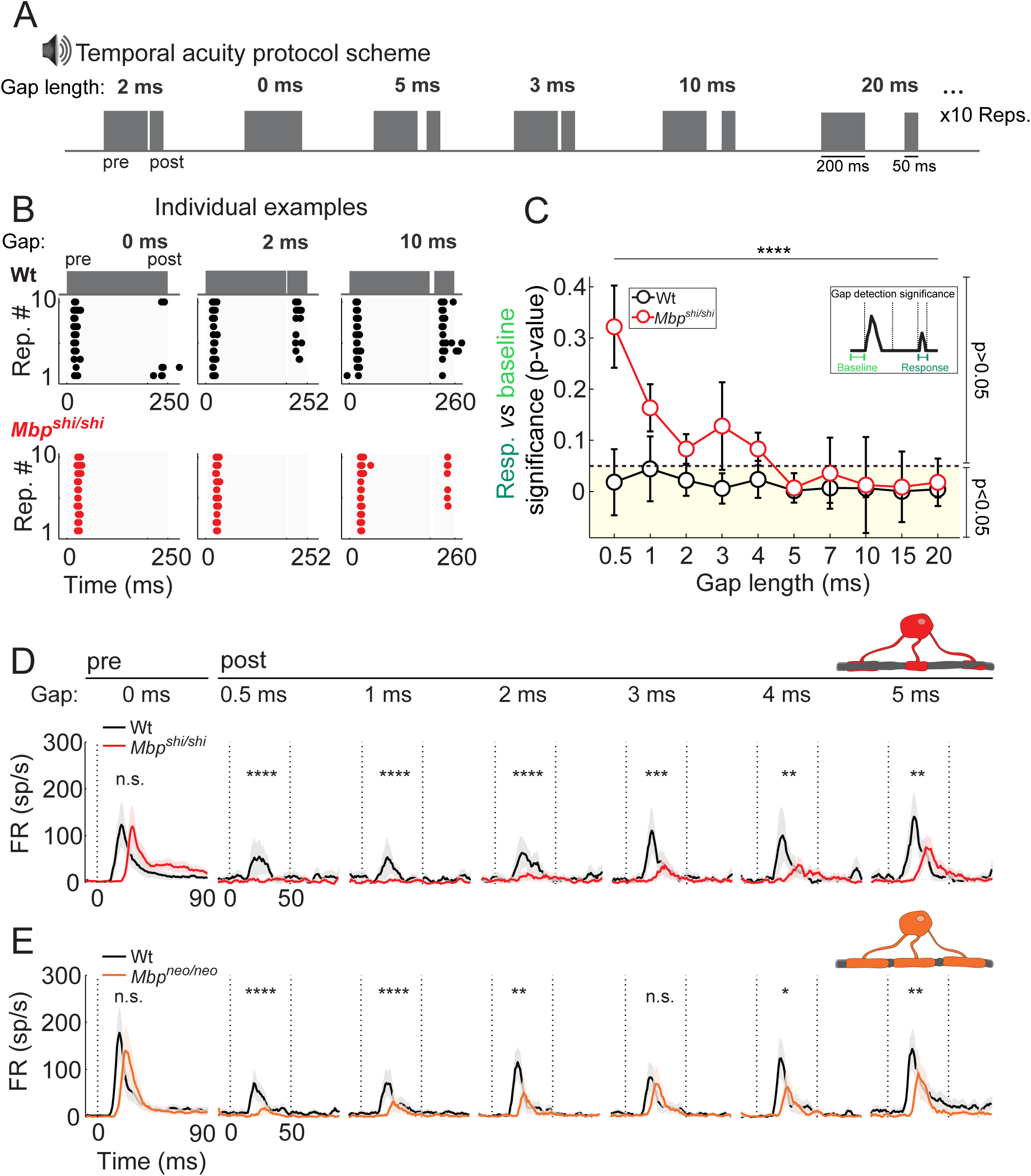
Cortical temporal acuity is impaired with CNS dysmyelination. **A)** Schematic of the gap-detection protocol used to test temporal acuity. A 200 ms long broad-band noise (BBN) pulse (pre) was followed by a silent gap and a second 50ms long BBN pulse (post). Gap lengths ranged from 0-50 ms. Each pre-gap-post sequence was repeated 10 times for every gap length. **B)** Individual example of ACx spikes (dots) evoked during sound presentation (gray patches) and gaps of 0 ms (left), 2 ms (center), and 10 ms (right) length in a Wt (upper, black) and a *Mbp*^*shi/shi*^ (lower, red) mouse during 10 stimulus repetitions (y axis) per gap length. **C)** Quantification of gap-detection significance (median statistical difference (p-value) between the baseline and the post-gap response per recording; see diagram above). A significant effect of gap-length between groups (p<0.0001), demonstrates that the gap detection rate in *Mbp*^*shi/shi*^ (n=14-20 recordings from 14 mice) is lower than for Wt (n=12-15 recordings from 12 animals) for the shorter gaps. Error bars: standard error of the median. Dotted line: threshold at 0.05. Yellow shadow: significant gap-detection. **D)** Average PSTH across animals for Wt (black; n=12) and *Mbp*^*shi/shi*^ (red; n=14) with S.E.M. as shaded area. Left to right: pre-gap responses followed by post-gap responses for 0.5 to 5 ms gaps. Significant effect of group was seen for all post-gap responses but not for pre-gap responses (p=0.46, p<0.0001, p<0.0001, p<0.0001, p=0.0002, p=0.0014, p=0.006; pre-gap and 0.5 to 5 ms post-gap respectively). **E)** Same as in D) but for *Mbp*^*neo/neo*^ (orange; n=8) mice and their respective Wt (black; n=7). Significant effect of group was seen for most post-gap responses but not for pre-gap responses (p=0.60, p<0.0001, p<0.0001, p=0.0015, p=0.19, p=0.01, p=0.003; pre-gap and 0.5 to 5 ms post-gap respectively). In D) and E), dotted vertical lines at time 0 ms depict the start of the sound, and at time 50 ms the end of the post BBN.

### Behavioral correlates of reduced temporal acuity

To understand the correlation between gap-detection deficits and perception at the behavioral level we performed two tests in behaving mice. First, we tested ‘gap-dependent pre-pulse inhibition of the acoustic startle reflex’ (GDIASR). This paradigm is widely used to behaviorally assess the detection of gaps within sound stimuli ^62–65^. For this test we could only use hypomyelinated *Mbp*^*neo/neo*^ mice that have no motor defects.

Pre-pulse inhibition of the acoustic startle reflex (ASR) requires sensorimotor gating. Acoustic startles are triggered by a loud unexpected sound ^66^ and can be measured by a piezo element under the animal (Figure S7A). A change in the sound background, for example a ‘silent gap’ just before the starling sound (Figure 6A), can inhibit the ASR if it is salient enough ^67^. In this paradigm, we used the level of inhibition by different ‘silent gap’ lengths (Figure S7B) as a behavioral measure of gap-detection (i.e. gap perception). Consistent with previous reports from wild type mice ^62,65,67,68^, shorter gaps elicited less inhibition than longer gaps (Figure 6B). Importantly, dysmyelinated *Mbp*^*neo/neo*^ mice showed a significant decrease in the inhibition of the ASR, when triggered by different gap lengths. Moreover, we determined a two-fold increase in the gap detection threshold (from 8 ms to 17 ms) (Figure 6B and 6C). Thus, deficits in temporal acuity, discovered at the physiological level in the ACx, were also paralleled at the perceptual level. We note that mice with slightly higher levels of *Mbp* expression, i.e. thinly myelinated heterozygous *Mbp*^*shi/+*^ mutants, did not show a reduction in basic startle or in gap-perception (Figures 6B, 6C, S7C and S7D). Thus, significant hypomyelination (>50%) is required to elicit gap-detection deficits at the behavioral level.

**Figure 6.**
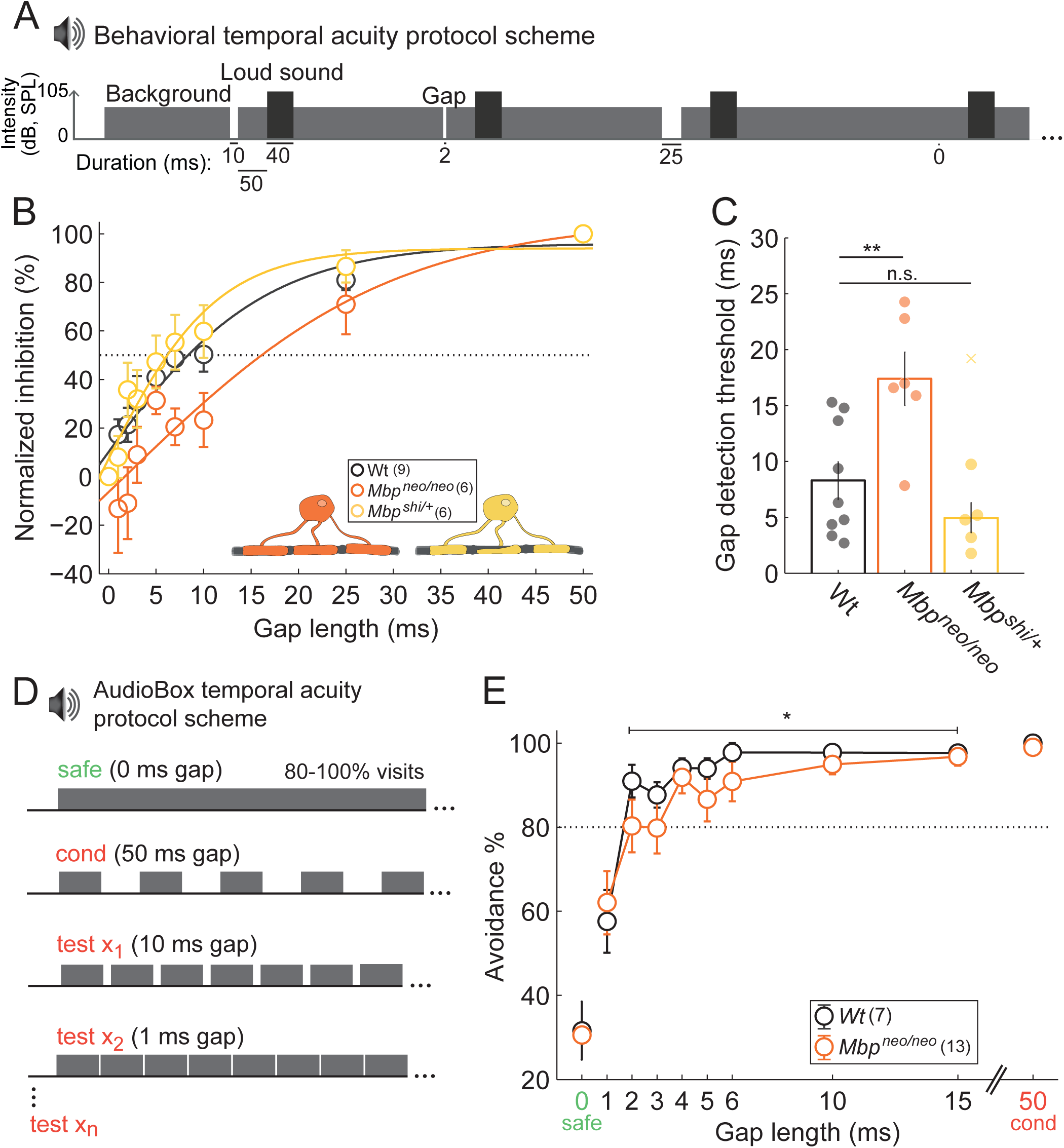
Behavioral temporal acuity is impaired by CNS dysmyelination. **A)** Schematic of the auditory startle reflex (ASR) sound protocol. Constant background broad-band noise (BBN; 70 dB) interrupted by a startle noise at random times (105 dB; 40 ms), occasionally preceded by a silent gap of varying length. All gaps presented were followed by 50 ms background sound before the startle appearance. Each gap-startle combination was repeated 10 times. **B)** The percentage of ASR inhibition elicited by the different gaps showed a strong relationship between the gap length and the startle inhibition. *Mbp*^*neo/neo*^ but not *Mbp*^*shi/+*^ mice (with a 50% reduction in *Mbp*) showed impaired inhibition of the ASR. Dotted line: threshold at 50% inhibition used for statistical analysis in C. **C)** Gap detection threshold is increased in *Mbp*^*neo/neo*^ mice (p=0.0048), but not *Mbp*^*shi/+*^ (p=0.36; outlier depicted with a cross), compared to Wt animals. **D)** Schematic of sounds used in the Audiobox paradigm. Safe visits to corner were accompanied by the safe sound (0 ms gap embedded in continuous BBN, ∼70 dB), and water was available. Conditioned visits were accompanied by the conditioned sound (cond; BBN with 50 ms gaps) water was not available and air-puff was delivered upon nose-poke. Once animals discriminated safe and conditioned, test visits with test sounds (gaps > 0 ms and <50 ms) were introduced. Water was available in these visits. **E)** Quantification of avoidance behavior (proportion of visits without nose-pokes) revealed a reduction (p=0.030) in *Mbp*^*neo/neo*^ animals (orange; n=13) for gaps >1 ms compared to Wt (black; n=7) confirming a behavioral deficit in temporal acuity in a naturalistic environment. All graphs show the mean per group. Error bars depict the S.E.M.

To confirm altered gap perception in freely behaving mice, we tested *Mbp*^*neo/neo*^ mutants in the ‘AudioBox’ (New*Behavior*, TSE systems), an automated system for auditory behavioral testing ^69–73^. Animals were exposed to ‘safe visits’ of one cage corner (Figure S7E), in which a continuous sound was played, and water was always available. However, during ‘conditioned visits’ of that corner the continuous sound was interrupted by a series of 50 ms ‘silent gaps’ that were paired (i.e. negatively conditioned) with short air-puffs to the nose when animals attempted to drink. We then compared the sensitivity to different gap lengths in mutants and controls ranging between 1 and 45 ms (Figures 6D and S7F). For each gap length we quantified the percentage of visits in which the mice avoided penalized nose-pokes, indicating perception of this gap. Despite equivalent sound exposure in the Audiobox (Figure S7G), gap detection threshold was increased to 4 ms in *Mbp*^*neo/neo*^ compared to the 2 ms threshold of wild type mice (Figure 6E). We conclude that in freely moving animals, myelination contributes to auditory perceptual acuity.

## Discussion

To explore the role of CNS myelination in information processing, i.e. beyond the known function of myelin in speeding axonal conduction, we studied the detection and perception of auditory stimuli in mice. We used auditory brainstem potentials and multiunit recordings in the auditory cortex of mutant mice, as well as behavioral readouts in awake animals, to compare the effect of partial (*Mbp*^*neo/neo*^) and complete (*Mbp*^*shi/shi*^) dysmyelination with that of an isolated reduction of glial metabolic support (MCT1^*+/-*^). The combined pattern of results reveals that diminished glial support of axonal metabolism, in the absence of dysmyelination, impairs sustained stimulus detection relevant for speech processing. The deficits were largely comparable, although partially discriminable from those resulting from dysmyelination. Oligodendrocytes, therefore, play an important role in ongoing perception through energetic support (Figure 7A), most evident when dysmyelination results in misdistribution of axonal ion channels (Figure 7B), and independent of a role in saltatory conduction. (Figure 7C).

**Figure 7.**
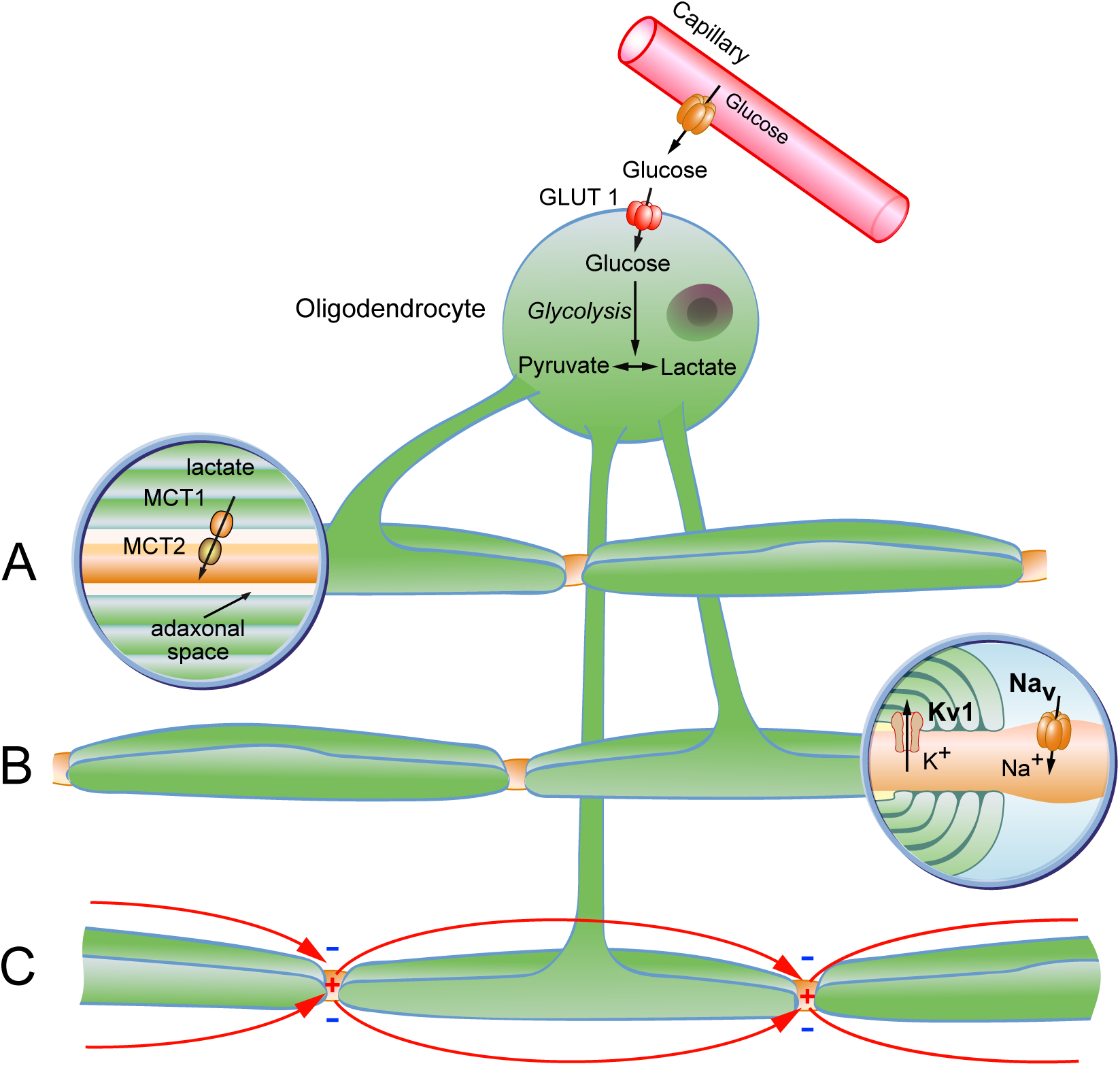
Role of oligodendrocytes in information processing extends beyond conduction velocity regulation to energy support of axons and excitability of axon initial segment and nodal. Schematic of oligodendrocyte and compact myelin (green) wrapping around 3 axons (brown) **A)** Glycolytic oligodendrocytes can support axons via the transport of glycolytic end products, such as lactate or pyruvate. **B)** The maintenance of axonal-excitable domains through clustering of ion channels depends on myelin compaction. **C)** Myelin is essential for the formation of axonal excitable domains and contributes to the speeding up of action potential conduction.

On the other hand, subcortical delays in dysmyelinated mice were accompanied by reduced synchrony of auditory brainstem potentials, a feature not observed in mice with only reduced glial metabolic support. The decreased amplitudes and shapes of waves II and III, were likely reflecting some desynchronization of the converging input at the level of the cochlear nucleus and olivary complex (where ascending auditory signals become bilateral). The later waves IV and V showed instead an increase in size in dysmyelinated mutants, most likely reflecting a compensatory mechanism to the loss of synchrony at the earlier stations. Similarly gain increases have been reported in mice deprived of auditory stimuli ^74^, and in humans as a result of aging ^44^ or multiple sclerosis lesions ^7,75–78^.

While dysmyelination had a strong effect on conduction velocity, spectral processing of simple stimuli (pure tones) in the primary auditory cortex appeared normal. Auditory information can reach the IC and then the ACx through different parallel pathways, each of which might be differently affected by dysmyelination. While simple click sounds, and pure tones are likely activating a limited (possibly the most direct) subset of sound encoding circuits, responses to these stimuli were normal, indicating that basic relay processing and sound integration is not affected by the absence of myelin. However, we detected deficits in temporal processing in a manner unexplained by reduced conduction velocity alone. Temporal reliability and acuity were affected in all mutants. Unexpectedly, the magnitude of this defect was correlated with the degree of hypomyelination but was not limited to dysmyelination phenotypes. It was also a feature of MCT1^*+/-*^ mice with an isolated metabolic support defect. Behavioral tests demonstrated that the temporal processing deficits, as detected by cortical multi-unit recordings, were paralleled at the perceptual level. Myelinating oligodendrocytes are thus clearly involved in the temporal processing of auditory signals, which have previously been implicated in higher CNS functions in humans, such as speech recognition ^79–81^. Here, the phenotypical similarities of two different mouse mutants, *Mbp*^*neo/neo*^ and MCT1^*+/-*^, suggest that compromised metabolic support alone is a critical component of reduced myelination and becomes detectable at the network level in the absence of any sign of axonal degeneration.

Myelination defects clearly affected the coding of a stimulus’ temporal envelope, reflected for example as ‘fatigability’, i.e. a lack of temporal reliability. For *Mbp* mutants it was unexpected that the responses to longer trains of clicks diminished or even stopped after 5 to 6 repetitions of the stimulus, especially when the intervals between consecutive clicks were 200 ms apart (repetition rate 5 Hz). Here, the altered distribution of sodium and potassium channels in unmyelinated regions is critical. In *Mbp*^*shi/shi*^ mice sodium channels are increased along bare axons ^82,83^, and abnormally high expression of Na_v_1.2 channels does not revert to Na_v_1.6 in development ^84,85^. This might contribute to increased fatigability in *Mbp*^*shi/shi*^ mice since Na_v_1.6 channels are not only associated with - but also necessary for - repetitive firing ^86^. In addition, an elongation of nodes ^58,87^ and overexpression of K_v_1.1 and K_v_1.2 potassium channels has been reported in *shiverer* mice ^58,88^. K_v_1 channels are normally clustered at juxtaparanodal domains ^89^ but may become abnormally exposed to the extracellular space upon dysmyelination and redistribute into the paranodal and internodal membrane causing increased K_v_1-mediated potassium efflux. In the absence of myelin, oligodendrocyte-dependent potassium siphoning would be reduced ^90^. Together, this might lead to a greater activity-driven accumulation of extracellular potassium, decreasing its driving force and resulting in a longer time course of membrane hyperpolarization. In agreement with these findings, when potassium channels in *shiverer* mice were acutely blocked with 4-AP in the *ex vivo* electrophysiological recordings of *Mbp*^*shi/shi*^ optic nerves (our accessible model of a dysmyelinated tract), we noted a partial restoration of single compound action potential (CAP) amplitude and a smaller hyperpolarization phase. The latter is consistent with the efficiency of 4-AP as a symptomatic treatment for MS ^91^.

Interestingly, reduced reliability was also observed in the MCT1^*+/-*^ mice, without dysmyelination, suggesting that reduced metabolic support to axons might be one of the responsible mechanisms. An indirect effect, and not necessarily an alternative explanation to the channel misdistribution, is the burden of dysmyelination and increased ion fluxes on the energy consumption of axonal Na^+^/K^+^ ATPases, postulated to be a contributing factor to axonal degeneration in demyelinated plaques of multiple sclerosis (MS) ^92^. Indeed, hearing difficulties have been reported also in MS patients ^78^. In the *Mbp* mutants, inadequate support might occur locally but have a dominant effect on the entire pathway. For example, lack of myelin might reduce energy reserves at higher spiking rates when the ATP-dependent ion pumps are most active. This might cause conduction blocks in the absence of axonal degeneration.

In addition to some loss of ‘temporal reliability’, we determined a myelination dependent defect of ‘temporal acuity’, i.e. a defect in the coding of rapid changes in the temporal structure of the stimulus. Here, a powerful paradigm is sound gap-detection ^54,93^, a function that relies on cortical auditory processing ^65,94–96^. In humans, defects in such a gap detection test are closely associated with a corresponding loss of accurate speech discrimination ^97,98^. Cortical neurons of dysmyelinated *shiverer* mice failed to detect silent gaps within a stream of white noise when these gaps were shorter than 3 ms, a marked deviation from control mice that detect gaps as short as 0.5-1 ms. Hypomyelinated *Mbp*^*neo/neo*^ mice exhibited a milder deficit.

Poor gap detection in *Mbp*^*shi/shi*^ mice could be caused by longer refractory periods and reduced excitability of the axons secondary to the abnormal distribution of ion channels. It would explain why these deficits in temporal acuity were less severe in the hypomyelinated *Mbp*^*neo/neo*^ mice. In contrast, neuronal ‘fatigue’ and the loss of temporal reliability are both caused by an underlying metabolic problem that is also prominent in MCT1 heterozygotes.

All multi-unit recordings were performed in anaesthetized mice. We note that key aspects of auditory processing occur ‘pre-attentively’ and should be unaffected by anesthetics ^99^. However, neuronal circuits with inhibitory input can behave differently under anesthesia, which can bias experimental results. Thus, it was important to parallel the electrophysiological results with behavioral experiments on auditory perception in freely moving animals. Since motor impaired *Mbp*^*shi/shi*^ mice could not be used for these experiments, we analyzed mice homozygous for the newly created *Mbp*^*neo/neo*^ allele, which had no visible motor defects and were long-lived. In this mutant, we confirmed poor gap detection at the perceptual level by operant conditioning of auditory stimuli and responses. We observed no such auditory deficits in heterozygous *shiverer* mice, which exhibit a minor dysmyelination ^36^. This suggests that a threshold level of hypomyelination is required for the auditory phenotype.

It has become increasingly clear that the role of myelination is not limited to speeding axonal conduction velocity. Moreover, the patchy myelination pattern of cortical circuits points to a broader role of oligodendrocytes and myelin in information processing ^100^. Our mouse models with predominant structural or metabolic defects of CNS myelin allowed exploring the role of myelination at the network level. This enabled us to mechanistically dissect reliability defects from acuity deficits in auditory processing. Thus, our studies revealed a novel role of myelin forming oligodendrocytes in information processing that goes beyond the speeding of neuronal responses. We discovered that myelin influences the sustained and temporally precise axonal firing, essential to properly code auditory stimuli. Moreover, specific phenotypical similarities between mutants with dysmyelination and metabolic defects suggests that a myelin-dependent energetic failure marks a common mechanism underlying auditory dysfunctions. Auditory phenotyping emerges as a promising experimental system to investigate the role of oligodendrocytes and myelination in information processing.

## Methods

### Mice

All mice were housed in standard plastic cages with 1-5 littermates in a 12 h/12 h light/dark cycle (5:30 am/5:30 pm) in a temperature-controlled room (∼21°C), with *ad libitum* access to food and water. All mice used were bred under the C57BL6/N background. Data obtained from male and female mice were pooled unless otherwise stated. The experimental and surgical procedures were approved and performed in accordance with the Niedersä chsisches Landesamt für Verbraucherschutz und Lebensmittelsicherheit (licence number 33.19-42502-04-16/2337).

Homozygous *shiverer* mice were obtained by crossing heterozygotes (*Mbp*^*shi/+*^). In all experiments, *Mbp*^*shi/shi*^ mice and their control littermates (Wt) were 6-12 weeks of age, unless otherwise stated.

A new hypomyelinated mouse with reduced *Mbp* expression (<50%), was generated by homologous recombination in ES cells, using a modified *Mbp* gene carrying a LacZ-neomycin cassette upstream of exon 1. Successful targeting disrupted the 5’ regulatory region without affecting the larger GOLLI transcription unit (Figure S1A). Homozygous *Mbp*^*neo/neo*^ mice were born at the expected frequency. For all experiments, *Mbp*^*neo/neo*^ and controls (Wt) were used at age 9.5-14 weeks, unless otherwise stated.

MCT1^+/-^ mice, with reduced axo-glial metabolic support ^3^ were kindly provided by Pierre Magistretti (Lausanne) and analyzed at age 10-14 weeks.

### Acoustic stimulation

All sound stimuli were digitally synthesized and presented using Matlab (The Mathworks^®^, USA), at a sampling rate of 98 kHz and in a pseudorandom order. There were typically 10 repetitions of each stimulus, unless otherwise stated. The intensities were measured in decibel sound pressure levels (dB-SPL). Sounds used for electrophysiology were delivered by an USB audio interface (Octa-capture, Roland, USA), amplified with a Portable Ultrasonic Power Amplifier (Avisoft, Germany) and played in a free-field ultrasonic speaker (Ultrasonic Dynamic Speaker Vifa, Avisoft, Germay). Calibration of the testing apparatus was made using BBN and click sounds at different intensities with a Brüel & Kjaer (4939 1/4”) free field microphone, with Brüel & Kjaer amplifier (D4039, 2610, Denmark). All experiments were performed in a sound-attenuated and anechoic room.

### Auditory Brainstem Responses

Auditory brainstem responses (ABRs) were measured as described ^101^. Mice were anesthetized with an intraperitoneal injection of 250 mg/kg Avertin (mixture of 2,2,2-tribromoethyl alcohol 2%, Sigma-Aldrich, and tert-amyl-alcohol 2%, Merck), except for the Mbp^*neo/neo*^ line, which was treated as in Jung et al. (2015) ^102^ and anesthetized with ketamine (125 mg/kg) and xylazine (2.5 mg/kg) i.p. Temperature was kept at 36° via a heating pad (World Precision Instruments, ATC 1000). For the recordings, subdermal needles (BD Microlance, 30G ½”, 0.3×13mm) were placed at the vertex (active electrode), the left pinna (reference) and the back of the animal (active shielding). Click stimuli were ipsilaterally delivered from a speaker located ∼9 cm from the left ear. Square click waveforms (0.03 ms long) were presented at 20 Hz (50 ms inter-trial intervals (ITI)) at different intensities (0-80 dB). The difference in potentials was amplified 10,000 times using a custom-made amplifier. A National Instruments shielded I/O connector block (NI SCB-68), interfaced in a Matlab environment was used for data acquisition (50,000 Hz sampling rate). Recorded voltage traces were bandpass filtered offline (300 - 3000 Hz) using a Butterworth filter. Data was cut from the stimulus presentation onset (0 ms) in 12 ms windows. Trials with heart rate artifacts (wave shapes larger than ± 4.7-9.2 μV) were removed, and 1000 trial repetitions of each stimulus were used for data analysis. Hearing thresholds were determined as the lowest intensity that evoked a response that was 3 times the root mean square of the response at 0 dB within a window of 1 ms centered on wave I. In addition, this measurement was confirmed by visual inspection of the lowest intensity that evoked a response (compared to 0 dB) within 6 ms from stimulus onset. The identity of the waves was determined by visual selection of the maximum peak value corresponding to each of the five waves according to the typical reported position of each wave (1 ms apart). For the analysis, an ANOVA was used to assess the changes in latencies or amplitudes of each wave, comparing control to mutant animals.

### Electron microscopy

After transcardial perfusion fixation (4% formaldehyde and 2.5% glutaraldehyde in phosphate buffer), mouse brains were dissected, and sagittal or coronal vibratome (Leica VT1000S) slices of 200-300 µm thickness were prepared. A punch of the region of interest was taken, prepared for electron microscopy ^103^ and embedded in Epon. Ultrathin sections were prepared using a Leica Ultracut S ultramicrotome (Leica, Vienna, Austria) and imaged with a LEO912 electron microscope (Zeiss, Oberkochen, Germany) using a 2k on-axis CCD camera (TRS, Moorenweis, Germany). G-ratios (axonal diameter/fibre diameter) were determined by analyzing 100 axons per animal selected by systematic random sampling on 10 random images taken at a magnification of 8,000 x.

### Acute electrophysiology

Prior to surgery, mice were anesthetized and maintained as reported for the ABR procedure. Mice were placed on a stereotaxic apparatus using inverted ear bars to avoid damage to the ear canal (World Precision Instruments, Sarota, FL USA, 502063). A cut was made along the midline and the skull exposed and cleaned of adherent tissue with a scalpel and hydrogen peroxide. A metal screw (M1×1, Germany) was inserted into the right parietal cortex and used as ground. A metal post was glued on the skull frontal to lambda with dental cement (Unifast, TRAD), this allowed for removal of the ear bars. The muscle temporalis was detached from the skull and a 4×2 mm craniotomy was performed using a dental drill (World Precision Instruments, Omnidrill3, tip #7), following the contour delimited rostral and ventrally by the squamosal suture, dorsally by the temporal ridge and caudally by the lambdoid suture. The speaker was placed ∼13 cm from the right ear of the mouse. At the end of every experiment, mice were euthanized by anesthesia overdose. The brain was removed, and immersion fixed in 4% PFA. After 24h, the brain was washed with PBS 1x and stored in 30% sucrose (Merk, K43386951).

Recordings were performed with glass-covered tungsten electrodes (AlphaOmega, Germany) or platinum/tungsten glass coated electrodes (Thomas Recordings, Germany) with impedances between 1.5 and 2 MΩ. In 30% of the recordings, the electrode was stained with Dye I (dioctadecyl-tetramethylindocarbocyanine perchlorate, Aldrich, 468495) dissolved in absolute ethanol, before insertion, for later visualization of its position. After the craniotomy, a drop of saline was used to clean the surface of the brain. The electrode was inserted perpendicular to the surface of the primary auditory cortex ^104^ using a micromanipulator (Kopf, Inc., Germany). We recorded extracellular multi-unit (MUA) sound-evoked responses in layer 3/4 of the ACx of anesthetized mice. Responses were characterized by onset latency and shape. ACx recordings in *Mbp*^*shi/shi*^ mice were done at a depth of ∼405 μm (62 μm standard deviation (sd)) without differences in depth between control and mutant mice (p=0.71). In the *Mbp*^*neo/neo*^ line, depth of recording was on average ∼350 μm (60.5 μm sd) without differences between groups (p=0.520). In MCT1^+/-^ mice, recordings were on average at ∼380 μm depth (37 μm sd) (p=0.32).

Electrophysiological signals were acquired at 32 kHz sampling rate, pre-amplified (HS-36-Led, Neuralynx, USA) and sent to an acquisition board (Digital Lynx 4SX, Neuralynx, USA) to yield the raw signals, which were acquired using a bandpass filter (0.1 or 200 Hz to 9000 Hz) and stored for offline analysis. Recording and visualization of the data was made using the Cheetah Data Acquisition System software (Neuralynx, USA). For multi-unit analysis (MUA), spikes that are not attributed to a single neuron, the signals were high-pass filtered at 350 Hz. For spike detection, a threshold of 6 times the mean absolute deviation from the median of the filtered voltage traces was used. For the analysis, only recordings that had significant auditory evoked responses at any sound intensity compared to a 200 ms pre-sound baseline activity (paired t-test) were used.

To measure tonotopy, *pure tones* of variable frequencies and intensities were presented (24 tones with frequencies between 2 to 31 kHz, and intensities between 0 and 80 dB) (Figure 3I). Each tone was 30 ms long, had on/off ramps of 5 ms. Presentation rate was 2 Hz (500 ms inter-trial interval). Five repetitions of each frequency/intensity combination were presented in a random order. The analysis was performed on individual recording sites, although in many mice recordings were obtained from several site along the ACx.

For the reconstruction of the single-site tuning (Figure 3J), the spikes elicited by each frequency in a 200 ms window after stimulus onset were added. The tuning curves (TC) were smoothened using a zero-phase digital hamming filter with a window of 4 points. The smoothened TC were then used to calculate the best frequency (BF; the frequency that elicited the maximum number of spikes at 60 and 80 dB). In normalized tuning curves, activity was expressed as a function not of frequency but of the distance (in octaves) from the best frequency. Response thresholds were obtained by visual inspection (blinded as to the genotype) of a 200 ms window from stimulus onset. Tuning curve width both at the base and the half distance from the peak activity (Figure 3M), was calculated as the z-score of the sum of spikes over a 60 ms window from stimulus onset at all frequencies and intensities.

The *click rate* (CR) protocol (Figure 4A) was used to assess temporal reliability. Each was a 0.05 millisecond long positive step function. Bursts of ten clicks were presented every 6 s, each at a different rate (2 - 50 Hz). The first click of each burst was used to characterize latency, amplitude, reliability and jitter of auditory response (Figure 3 and S3). For the *latencies* measurement we only included the spikes that occurred at least 10 ms after sound onset, given latencies reported for the ACx in anesthetized mice ^104^. To measure reliability of repeated responses (click rate protocol, Figure 4), synchronicity measurements were obtained from a measure of the vector strength ^105^ of the spikes occurring for the duration of the stimulation in all repetitions of the stimulus was obtained by converting each spike time (τ_*i*_) as a circular vector with phase (θ_*i*_) between 0 and 2π, according to the stimulus phase of a specific rate. All the spikes occurring after each of the 10 clicks were selected in terms of their latency respective to the previous click. A window equivalent to the phase duration (equal to the inter-click interval) was selected for each click rate presentation. The spike phases were then expressed in terms of: 

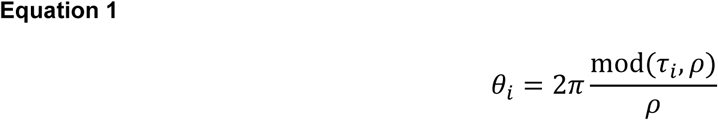

Here, τ_*i*_ is the latency of each spike, θ_*i*_ was the phase representation of each spike, and ρ was the phase of the stimulus presented (for example for a click train at 5 Hz, 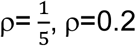). To calculate the percentage of spike synchronicity, the total spike phase distribution was binned in 20 bins from 0 to 2π, and the spike count of the bin with the maximum synchronization in the first half of the vector (π), was taken. This spike count was expressed in terms of percentage, which means the percentage of spikes that were fully synchronized per stimulus in one recording.

The *gap-in-noise detection* paradigm was used to evaluate temporal acuity. Once every second, 200 ms long broad band noise (BBN, pre-gap) sound was followed, after a short silent gap, by a 50 ms long BBN sound (post-gap). Silent gaps durations were 0, 0.5, 1, 2, 3, 4, 5, 7, 10, 20, 50 and 100 ms (Figure 5A). Rise/fall time of the BBN pulse was 1 ms. For *gap-detection* analysis, only recordings with a significant response to the mean of the pre-gap sound (sum of spikes over 100 ms window after stimulus onset), compared to pre-sound baseline, were used (paired t-test). For each animal, recordings at different locations were averaged. Additionally, only those animals that had a significant response to the post-gap BBN following a 100 ms gap were included in the analysis (100 ms window from the start of the post-gap BBN). Peri-stimulus time histograms (PSTH) were generated by adding spikes across trials in 1 ms bins over a 100 ms window (Figure 5D). The pre-gap response (PSTH in a 100 ms window) was generated from all gap trials, since the pre-gap response should not be affected by the gap that follows it. For the post-gap responses (PSTH over 100 ms), each gap was treated separately. Comparisons between control and *Mbp*^*shi/shi*^ and *Mbp*^*neo/neo*^ mice revealed no significant differences in the amplitude of the pre-gap response (ANOVA, p=0.53 and p=0.6 respectively). Also, the peak of the post-gap response was used for group comparisons. Since some groups showed increased latencies, the PSTHs were centered at their peaks. An ANOVA was performed in a 21 ms window centered on the peak.

For the analysis of the gap-detection *vs* baseline, a 50 ms window of baseline activity was compared to a 50 ms window of the post-gap response for each recording site using a paired t-test.

### Gap pre-pulse inhibition of the acoustic startle reflex

We used behavioral Gap-PPI to test behavioral temporal acuity ^67^. This test was performed in *Mbp*^*neo/neo*^, *Mbp*^*shi/+*^ and their control littermates. We used 9 control animals (wild types pulled together from both dysmyelination lines), 6 *Mbp*^*neo/neo*^, 6 *Mbp*^*shi/+*^. All mice were between 9 and 16 weeks at the age of testing. For testing, a mouse was confined in a custom-made Plexiglas tunnel (12 cm long by 4 cm diameter), placed above a piezo element (TRU components, 800 Ohms impedance, 50 mm diameter, spanning 30V) (Figure S7A). The piezo was connected to a data acquisition system (NI SCB-68). Data acquisition (1 kHz sampling rate) was controlled with a custom-made code in a Matlab and sound delivery was done using pre-synthetized sound tracks in Matlab, played through VLC media player (Video Lan Organization).

Testing protocol was as summarized in Figure S7B. Mice were initially acclimatized to the setup for 10 min before the start of the experiment: 5 min. of silence followed by 5 min. of exposure to the background sound (BBN of 70 dB). Testing was then initiated. The startle noise was a BBN 105 dB with 40 ms duration. It appeared every 10-20 s. at random times to avoid startle prediction. The startle could be preceded by a gap (0, 1, 2, 3, 5, 7, 10, 25 or 50 ms long) which ended 50 ms before the startle (Figure 6A). Gaps and startle had on/off ramps of 1 ms. The session started with 10 startle-only pulses, followed by gap preceded startle presentations (10 startle presentation per gap type). The experiment ended with five startle-only pulses, to assess habituation. The complete presentation of 105 trials had a duration of approximately 30 min.

*Gap-PPI* analysis was done as described before ^67^, with mild modifications. The magnitude of the acoustic startle reflex (ASR) was measured as the maximal vertical force (peak-to-peak voltage output) exerted on the piezo element in a 500 ms window starting with the onset of the startle noise. Baseline activity (2 times the root mean square of the voltage trace in a 500 ms window before the startle noise) was subtracted. Startle and baseline amplitude were measured in arbitrary voltage units provided by the piezo measurement. Noisy trials (3 times the standard deviation of the root mean square of a 500 ms window before gap presentation) were discarded from the analysis. The percentage of pre-pulse inhibition for each gap and mouse was calculated following: 

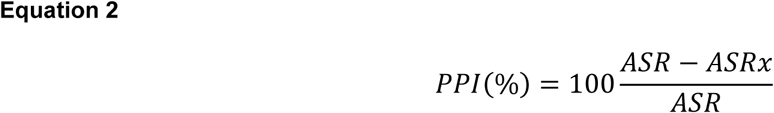

Where ASR is the startle response elicited at 0 ms gap, and ASRx is the startle response elicited per gap played. The data were fitted with a generalized logistic function: 

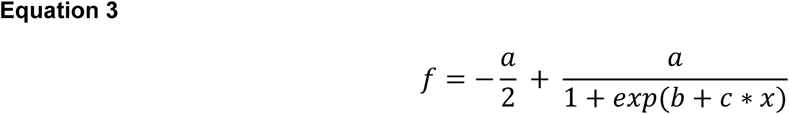

Recordings with a fit coefficient (R^2^) below 0.6 were excluded from the analysis.

The gap detection threshold was considered as the value of the fitted curve that elicited 50% of the maximal inhibition achieved per mouse. A second normalization was then made to the longest gap (50 ms) presumed to elicit the maximum value of inhibition (100%). For the statistical comparisons of the gap-detection threshold and amplitude of startle, parametric or non-parametric t-test were used. A 2-way ANOVA was performed for the comparison of the inhibition of the ASR along the different gap lengths.

### AudioBox

The AudioBox system was used to test behavioral gap-detection. The AudioBox (New*Behavior*, TSE systems) is an automated system for behavioral acoustic conditioning ^69–72^. This system is designed to minimize the interference from the experimenter and provides a spatially and socially enriched environment in which female mice live in group (up to 10 mice) for several days. The setup consisted of a home-cage, where animals had food *ad-libitum*, and a corner within a sound-attenuated box, where they could access water also *ad-libitum* (Figure S7E). A speaker was over the corner (22TAF/G, Seas Prestige). Animals carried a transponder (implanted under anesthesia, see ^72^) that was detected by an antenna at the entrance of the corner. Time spent in the corner, number of nose-pokes and sound played were detected through sensors in the corner.

For these experiments, *Mbp*^*neo/neo*^ mice were chosen over *Mbp*^*shi/shi*^ because of their normal life span and lack of motor impairments. Three experiment replications were pooled together. A final number of 7 control animals (Wt) and 13 mutant animals (*Mbp*^*neo/neo*^) that were 6-14 weeks old at the start of the experiment were used.

Behavioral testing occurred in the corner. Each time a mouse entered the corner a “visit” started, and a sound was played for the duration of the visit. The training paradigm, summarized in Figure S7F, consisted of 3 phases. During the habituation (one day doors opened, and 3-9 days doors closed), the safe sound (continuous BBN) was played in all visits. In this phase, animals could nose-poke and get access to water. During the conditioning phase ‘conditioned’ visits were introduced (Figure 6D and S7F). In safe visits, as before, the animal could make a nose-poke and get water. Animals that did not nose-poke in more than 60% of safe visits were immediately excluded from the behavioral experiment. This was the case for 11 out of a total of 18 Wt and 7 out of 20 *Mbp*^*neo/neo*^ tested. In conditioned visits (pseudo-randomly distributed according to probabilities as in Figure S7F), however, a conditioned sound (BBN interrupted by 50 ms silent gaps every 500 ms) was played, and a nose-poke triggered an air-puff, and no opening of the doors. Mice had to learn to not nose-poke in visits in which silent gaps interrupted the sound. Typically, this occurred 1-2 days after introduction of the conditioned visits. The percentage of conditioned sound presentation was increased stepwise 5% every 2 days. After the conditioning phase, the gap testing phase began. Every 4 days two new visit types were introduced, each with BBN sounds interrupted, like the conditioned sounds, by gaps, which varied in length with visit type (Figure 6D and S7F). During this phase 70% of visits were safe, 20% conditioned and 10% conditioned gaps. A total of 15 gaps were tested with lengths between 1 and 45 ms.

The new sounds were chosen in pairs in a semi-random fashion, such that the more difficult gaps (gaps below 5 ms) were presented in the same block of days as gaps longer than 5 ms. A minimum of four days with each pair of sounds was allowed before starting a new gap testing.

The percentage of avoidance was quantified as the number of visits without nose-pokes for each sound played, only considering the first two days of training for each sound. The number of visits was compared to ensure equivalent sound exposure. The data from visits with gaps between 2 and 15 ms duration were used for statistical comparison.

### Analysis generalities

Analyses and figure outputs were always performed in a Matlab environment. All confidence intervals correspond to α=0.05. Significance corresponds to ^n.s.^p>0.05, *p≤0.05, **p≤0.01, ***p≤0.001, ****p≤0.0001. Statistical analysis was parametric or non-parametric according to the Shapiro-Wilcoxon normality test.

Paired comparisons were done using t-test or ranksum and post-hoc multiple comparisons were performed using 2-way ANOVA or Kruskal-Wallis.

## Acknowledgements

The authors would like to thank Gerhard Hoch for the ABR amplifier and for technical support; Gudrun Fricke-Bode, Annette Fahrenholz, Saskia Heibeck, Uschi Müller, and Swati Subramanian for help with molecular and histological methods; Cornelia Casper for help with animal care; Chi Chen for assistance with the behavioral setups; Alexander Dieter for programming support; Jan Ficner for Figure 7 design; and Pierre Magistretti for providing MCT1 heterozygous mice. Funding was granted by the Max Planck Society and the cooperation program CONACyT-DAAD for S.M. (scholarship 381831), and an ERC Advanced Grant (K.A.N.). M.M. was funded through, K.A.N. and W.M. were supported by the Cluster of Excellence and DFG Research Center Nanoscale Microscopy and Molecular Physiology of the Brain.

## Author contributions

Conceptualization, L.dH. and S.M.; electron microscopy images, T.R. and W.M.; ABR recordings of *Mbp*^*neo/neo*^, N.S.; ABR recordings, *in vivo* electrophysiology, and behavior, S.M.; *in vivo* electrophysiology for SSA, L.d.H.; generation, cellular and molecular characterization of the *Mbp*^*neo/neo*^ model, W.M., and M.M.; AIS stainings and quantification, I.T., A.B., and M.K.; ON recordings, S.M., and A.T.; maintenance of MCT1 mice, K.K.; data analysis, S.M., and L.dH.; data curation, S.M.; writing-original draft, S.M., L.dH., and K.A.N; revision-editing, all authors; funding acquisition, S.M., L.dH., and K.A.N; project supervision, L.dH., and K.A.N.

