## Supplemental Information for "A role of oligodendrocytes in information processing independent of conduction velocity"

#### Methods

##### Acute electrophysiology

Response *amplitude* was the measured as the sum of the spiking activity across 10 repetitions of the first click stimuli of the 5, 8, 10, 14 and 20 Hz condition. *Reliability* measurements were obtained from the same data used for the amplitude. The percentage of trials that contained at least one spike within a 37 ms window that started 12 ms after stimulus onset was obtained. *Jitter* was measured also under these conditions as the standard deviation from the mean $1^{st}$-spike latency value. All measurements were performed in single recording sites and then averaged for each animal. All comparisons were done using parametric or non-parametric t-tests.

*Frequency oddball paradigms* were evaluated presenting two pure-tones ($f_1$ and $f_2$) separated in frequency by $\Delta f=0.1$ as shown by **Equation 1**

$$\Delta f=(f_2 - f_1)/( f_2 \times f_1)^{1/2}$$

$f_1$ and $f_2$ were centered on the best frequency of the MUA at each recording site. Three different probabilities of presentation of the deviant sound were used (5%, 10% and 20% of appearance, at 3 Hz). Each combination of stimuli (Δf, % of appearance and rate) was presented 300-500 times.

The SSA index (SI) was calculated as described in [106]:

$$SI(f_i) = (d(f_i) - s(f_i)) / (d(f_i) + s(f_i))$$

where $_i$ = 1 or 2 and $d(f_i)$ and $s(f_i)$ are responses (as normalized spike counts) to frequency $f_i$ when it was deviant or standard, respectively. With increasing Δf (e.g. 20%) we observe a greater SI, since the response to the deviant is larger due to the greater difference between $f_1$ and $f_2$. This happens because similar frequencies are generalized and are susceptible to adaptation. A small Δf (e.g. 5%) will generate a reduced SI. Changing the presentation rate also modifies the SI, since very low rates of presentation (e.g. 1 Hz) will not generate a continuity of the sound and the sounds will be computed as random events. The higher the presentation rate (e.g. 3 Hz), the easier to detect deviant stimuli and the larger the SI. In addition, the probability of deviant presentation also modifies SI. If a deviant sound is very rare (e.g. dP=5%) its detection will be easier than if the sound occurs more frequently (e.g. dP=20%).

The *SSA index* is calculated as follows:

**Equation 6**

$$SSAi = \frac{\partial s - \rho s}{\partial s + \rho s}$$

Where $\delta s$ represents the deviant sound and $\rho s$ the standard sound.

**Histology and microscopy**

Before cutting, brains were placed in PBS 1x for 24 h. Coronal sections (100 µm thickness) were obtained using a vibratome (Leica VT1000). Individual slices were incubated in PBS 1x with DAPI (Sigma-Aldrich) 1:40,000 for 5 min/25 rpm at room temperature. After incubation, the slices were washed twice with PBS 1x, 5 min/25 rpm. The slices were mounted into microscope slides (Menzel-Gläser, 76x26 mm), dried and covered with mounting media (Aqua Polymount, Polysciences Inc, 18006) and cover slips (Menzel-Gläser, 24x50 mm, #1). The slides were stored in dark at ~4°C for further visualization with a confocal microscope (Zeiss 510 meta).

**AIS stainings**

**Solutions and antibodies**

Cutting solution contained (in mM): 125 NaCl, 25 $NaHCO_3$, 1.25 $NaH_2PO_4$•$H_2O$, 3 KCl, 25 glucose, 1 $CaCl_2$, 6 $MgCl_2$, 1 kynurenic acid Blocking solution contained 5% goat serum and 0.5% Triton X-100 in PBS. Primary antibodies used were Anti-$K_V$7.3 (KCNQ3n) raised in guinea pig (1:200) kindly provided by Ed Cooper (Baylor College of Medicine, Houston, TX, USA) (Jin et al., 2009). AnkyrinG antibody was raised in rabbit (1:100; Santa Cruz). Secondary antibodies: Alexa 488-conjugated donkey anti-rabbit (1:1000, Invitrogen) and Cy3 donkey anti-guinea pig, (1:1000, Dianova).

**Immunostaining**

Mice were euthanized by cervical dislocation, the brain removed, submerged in ice-cold cutting solution and coronal 300 µm slices were prepared using a vibratome (VT1000S, Leica) with cutting amplitude of 1 and 0.05-0.07 forward speed. Freshly cut slices were fixed with Methanol chilled at -20°C after removal of the cutting solution and incubated at -20°C for 10 min. The slices were washed with PBS 1X (3 times, 5 min. each) and blocked for 2.5-5 hours at room temperature. The primary antibodies dissolved were added after blocking and slices incubated for 2 days at room temperature. Slices were washed with PBS 1X (3 times, 5 min. each), and secondary antibodies were added and left for 2 hours at room temperature (protected from the light). The slices were washed with PBS 1X (4 times, 5 min. each), stained with DAPI (4',6-diamidino-2-phenylindole) for 10min., and then washed with PBS 1X (4 times, 5min. each). Slices were mounted using Aqua-Poly/Mount (Polysciences, Inc) and stored at 4°C protected from the light.

AIS were imaged using a confocal microscope (Zeiss 510 meta). Z-stacks were acquired using a 40x objective (1 µm step, 30 z per stack in average). 5 fields were imaged per mouse.

**Analysis**

AIS length analysis was performed from confocal scans. The immuno-signal of AnkyrinG and $K_V7$ were measured with the segmented line tool in FIJI (v.1.51) by drawing a line along the AIS. For both channels the beginning and end of the expression was defined when the immuno- signal was stronger than the background. No background subtraction was applied. Comparisons were done for all AIS quantified per animal and using a t-test for comparison between *shiverer* mutants and wild-type mice.

**Optic nerve recordings**

**Solutions**

Artificial cerebrospinal fluid (aCSF) was prepared, containing (in mM): 124 NaCl, 3 KCl, 2 $CaCl_2$, 2 $MgSO_4$, 1.25 $NaH_2PO_4$, 23 $NaHCO_3$ and 10 glucose monohydrate. The solution had a pH of 7.4 and was controlled for osmolarity.

**Electrodes**

Both stimulating and recording suction electrodes were fabricated from borosilicate glass capillaries as previously described [60]. An Ag/AgCl wire was inserted into the electrode, and the capillary space back-filled with aCSF with 10 mM glucose.

**Optic nerve preparation and stimulation**

Experiments were done as reported before [4,5]. Mice were decapitated and the skin over the skull was removed. The optic nerves were separated from the eyes at the ocular cavity. Then, the skull was opened, and the optic nerves detached by cutting caudally to the optic chiasm. The preparation was placed into an interface perfusion chamber (Harvard Apparatus, Holliston, MA) and continuously superfused with carbogen (95% $O_2$, 5% $CO_2$) bubbled aCSF at 37°C during the experiment at about 4 mL/min. Optic nerves were detached from the chiasm and placed in the suction electrodes for recording. The stimulation electrode was attached to the end that was near the retina. Continuous stimulation of 0.75 mA at 0.1 Hz was performed until the recorded compound action potential (CAP) showed a steady shape (typically between 10 and 40 min.). For stimulation, the electrode was connected to a battery (Stimulus Isolator 385; WPI, Berlin, Germany) that delivered a supramaximal stimulus to the nerve. The recording electrode was connected to an EPC9 amplifier (Heka Elektronik, Lambrecht/Pfalz, Germany). Signals were amplified 500 times, filtered at 30 kHz, and acquired at 100 kHz.

**Analysis**

The CAP consists typically of 3 peaks of different amplitudes and latencies. Peak 1 is the fastest (approximately at 0.5 ms after stimulation) and it reflects the activity of high caliber myelinated axons Peak 2, at 1ms, reflects the activity of middle caliber axons, and peak 3 (1.5 ms) of small caliber axons. Since in *Mbp$^{shi/shi}$* mice it was not possible to distinguish peak 1, the analysis was done independently of peak specification. We measured the maximal amplitude elicited by specific stimulation and the correlated area under the curve. The CAP area is a reliable measure that reflects the proportion of axons that are activated by the stimulation [4,5,60]. For these recordings, 6 to 10 nerves from 3 to 9 animals per group were used and pooled in the analysis.

CAP amplitude was measured as the maximum data point of the whole CAP curve. We believe the measurements mainly reflect the activity of middle caliber axons. For the detection of the depolarization amplitude, a window of 3.7 ms after the stimulation artifact was taken, to ensure detection of peaks even in the mouse models with delayed responses. For the quantification of the hyperpolarizing peak, a window of 3 ms around the maximum negative peak was taken. Statistical analysis of depolarization and hyperpolarization peaks was done using 2-way ANOVAs.

For the measurement of conduction velocity, we obtained the latency to the maximum peak at a stimulation of 0.7 mA, and the length of the nerve stretched between the electrodes, which was measured after the end of every experiment. The latency/length value was normalized to the mean of conduction velocity values of the control animals. Parametric t-test was performed for statistical analysis.

### Figure S1

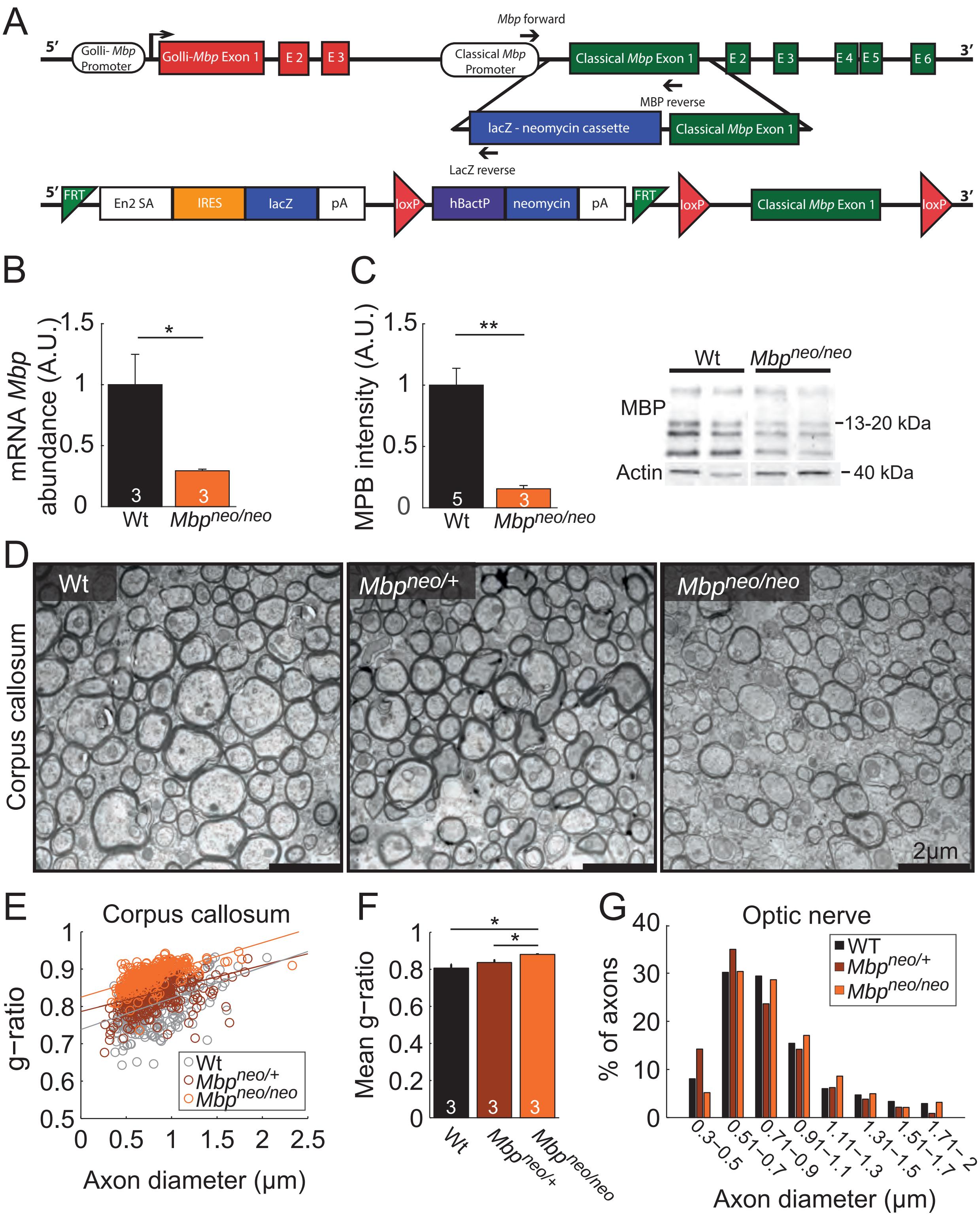
A
5′
Golli- Mbp Promoter
Golli-Mbp Exon 1
E 2
E 3
Mbp forward
Classical Mbp Promoter
Classical Mbp Exon 1
E 2
E 3
E 4
E 5
E 6
3′
MBP reverse
lacZ - neomycin cassette
Classical Mbp Exon 1
LacZ reverse
5′
FRT
En2 SA
IRES
lacZ
pA
loxP
hBactP
neomycin
pA
FRT
loxP
Classical Mbp Exon 1
loxP
3′
B
mRNA Mbp abundance (A.U.)
1.5
1
0.5
0
*
3
3
Wt
Mbpneo/neo
C
MPB intensity (A.U.)
1.5
1
0.5
0
**
5
3
Wt
Mbpneo/neo
Wt
Mbpneo/neo
MBP
13-20 kDa
Actin
40 kDa
D
Corpus callosum
Wt
Mbpneo/+
Mbpneo/neo
2μm
E
Corpus callosum
g−ratio
1
0.9
0.8
0.7
0.6
0.5
0
0.5
1
1.5
2
2.5
Axon diameter (μm)
Wt
Mbpneo/+
Mbpneo/neo
F
Mean g−ratio
1
0.8
0.6
0.4
0.2
0
*
*
3
3
3
Wt
Mbpneo/+
Mbpneo/neo
G
Optic nerve
% of axons
40
30
20
10
0
WT
Mbpneo/+
Mbpneo/neo
0.3−0.5
0.51−0.7
0.71−0.9
0.91−1.1
1.11−1.3
1.31−1.5
1.51−1.7
1.71−2
Axon diameter (μm)

**Figure S1. Molecular characterization of *Mbp*$^{neo/neo}$ model**

**A)** Genetic construct including the lacZ-neo cassette as provided by the European Conditional Mouse Mutagenesis Program (Eucomm). Eucomm consortium website with official depiction of construct: http://www.mousephenotype.org/about-ikmc/eucomm-program/eucomm-targeting-strategies. **B)** *Mbp* expression at the mRNA level and **C)** MBP expression at the protein level showed a significant reduction in *Mbo*$^{neo/neo}$ mice (~70%, p=0.047 for mRNA; and 80%, p=0.0037 for MBP). Inset shows Western Blot example used for quantification. **D)** Electron microscopy images of Wt (left) *Mbp*$^{neo/+}$ (middle) and *Mbp*$^{neo/neo}$ (right) caudal corpus callosum transverse sections. *Mbp*$^{neo/neo}$ mice have thinner compacted myelin than Wt. Scale bars: 2 µm. **E)** Distribution of g-ratios (axon diameter/axon + myelin diameter) per axon caliber in Wt (gray), *Mbp*$^{neo/+}$ (brown) and *Mbp*$^{neo/neo}$ (orange) caudal corpus callosum. **F)** G-ratio quantification revealed that *Mbp*$^{neo/neo}$ mice have significantly thinner myelin than Wt (t-test, p=0.022) and their heterozygote littermates *Mbp*$^{neo/+}$ (t-test, p=0.023). No differences between Wt and *Mbp*$^{neo/+}$ (t-test, p=0.27) **G)** Axon caliber distribution was not changed in *Mbp*$^{neo/neo}$ animals.

### Figure S2

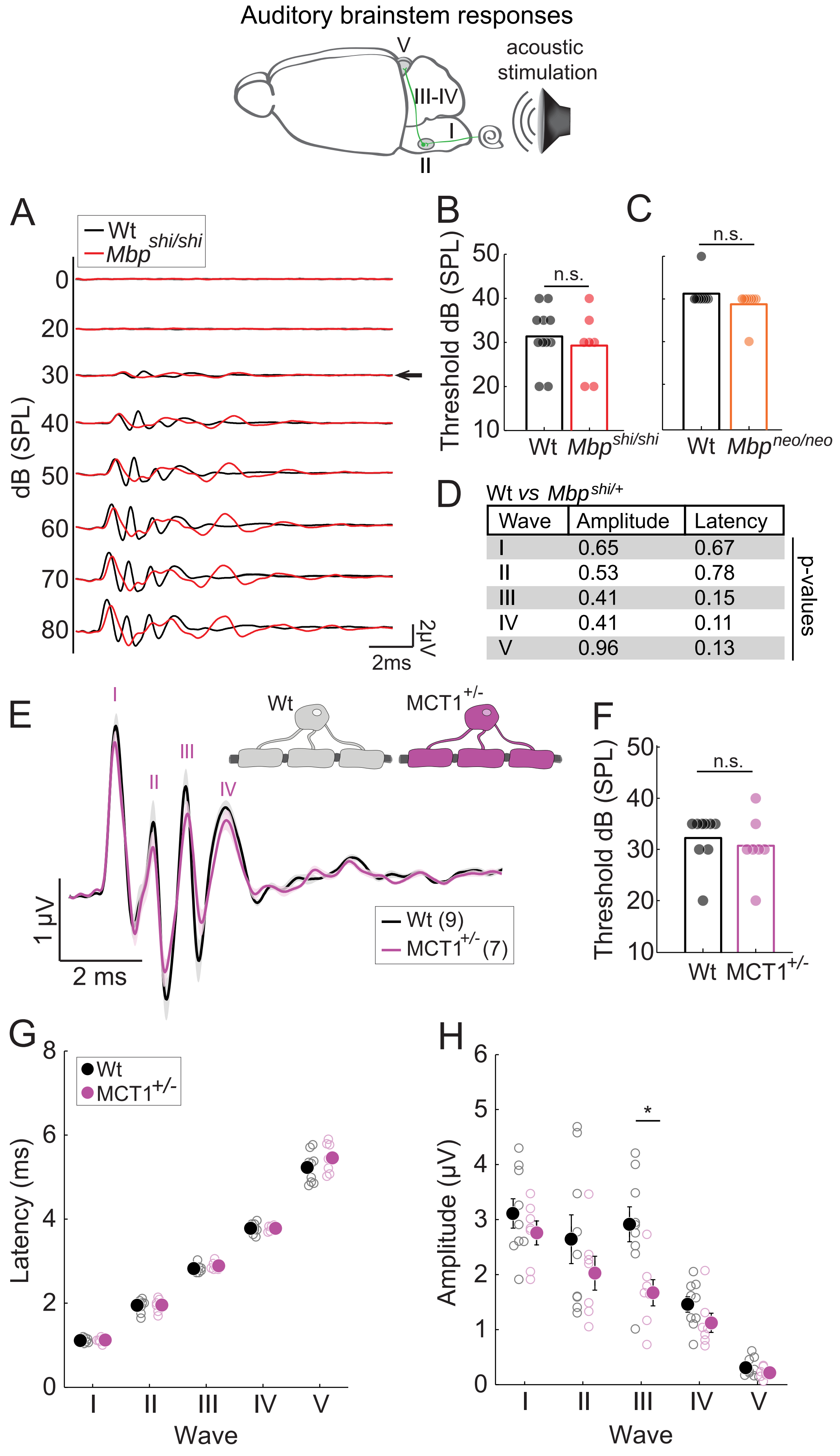

Auditory brainstem responses
acoustic stimulation
V
III-IV
I
II
A
Wt
Mbp shi/shi
dB (SPL)
0
20
30
40
50
60
70
80
2μV
2ms
B
Threshold dB (SPL)
n.s.
Wt
Mbp shi/shi
C
n.s.
Wt
Mbp neo/neo
D
Wt vs Mbp shi/+
Wave Amplitude Latency
I 0.65 0.67
II 0.53 0.78
III 0.41 0.15
IV 0.41 0.11
V 0.96 0.13
p-values
E
Wt
MCT1+/-
1 μV
2 ms
Wt (9)
MCT1+/- (7)
F
Threshold dB (SPL)
n.s.
Wt
MCT1+/-
G
Wt
MCT1+/-
Latency (ms)
Wave
H
*
Amplitude (μV)
Wave

**Figure S2. ABRs thresholds are affected neither by CNS dysmyelination nor by a reduction in glial-metabolic support.**

Schematic of circuit origin of the different auditory brainstem response (ABR) peaks. **A)** Mean across animals of ABRs to clicks at different intensities for control (black; n=11) and *Mbp*$^{shi/shi}$ mice (red; n=7). **B)** Quantification of thresholds, intensity at which first significant response is observed, revealed no significant difference between control and *Mbp*$^{shi/shi}$ mice (p=0.54). **C)** Mbp$^{neo/neo}$ (orange; n=8) thresholds also did not differ (p=0.53) from wild type littermates (black; n=8). **D)** No difference between the 7 *Mbp*$^{+/+}$ and 4 *Mbp*$^{shi/+}$ mice combined as controls for the *Mbp*$^{shi/shi}$ group as illustrated by table of p-values for amplitude and latency for individual peaks at 80 dB. **E)** Mean ABR response elicited by clicks at 80 dB in wild type (black, n=9) and MCT1$^{+/-}$ mice (purple, n=7) were comparable in shape and latency. **F)** Auditory thresholds were not different between groups (p=0.39). **G)** Latency of waves I-V elicited by clicks at 80 dB was comparable between groups (p=0.84, p=0.92, p=0.18, p=0.97, p=0.23, respectively). **H)** Amplitude of waves I-V, revealed only a mild decrease in wave III for MCT1$^{+/-}$ mice (p=0.34, p=0.29, p=0.01, p=0.15, p=0.22 for waves I to V respectively). All measurements depict the mean of the recorded animals and error-bars (or shaded errors) correspond to the S.E.M.

### Figure S3

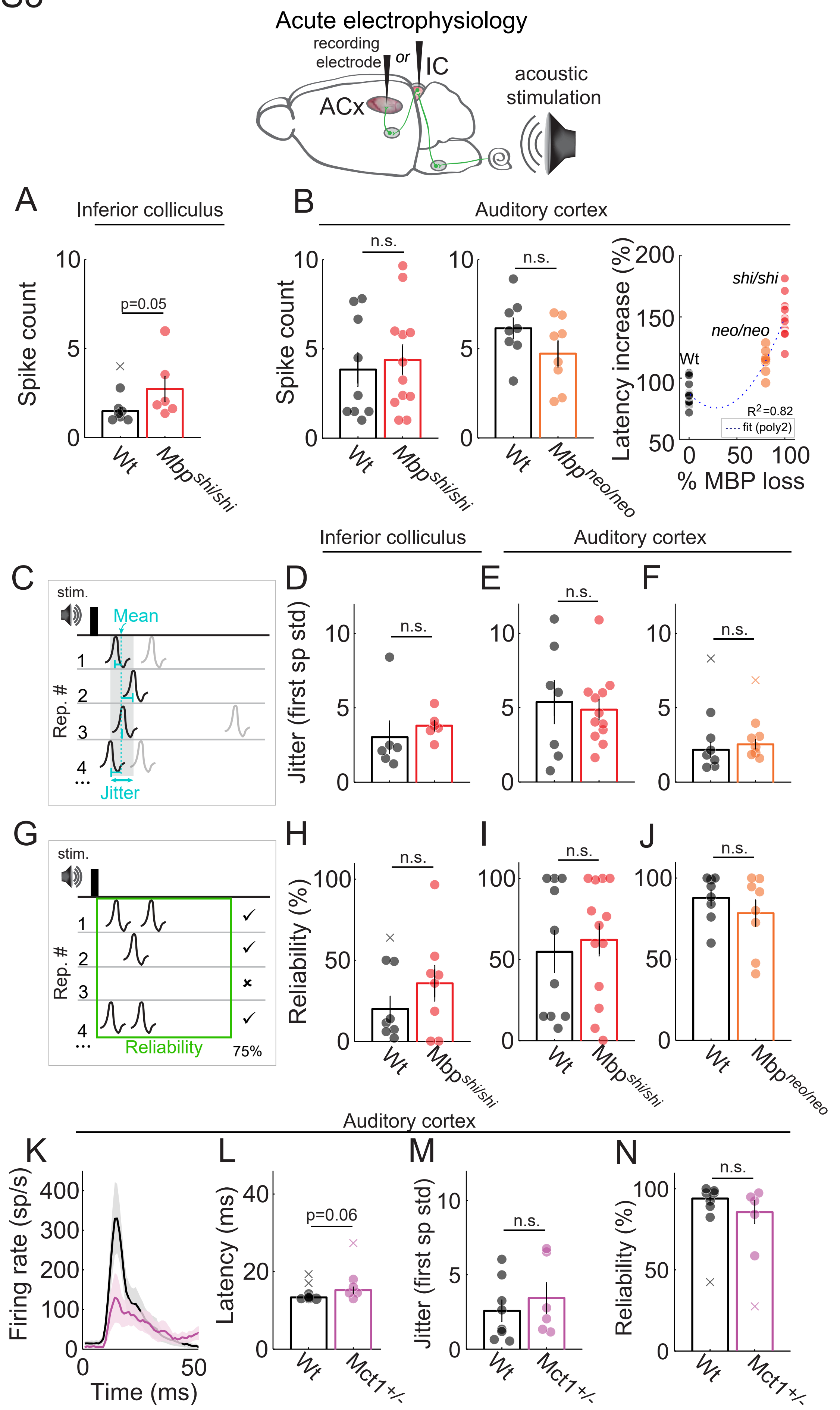
Acute electrophysiology
recording electrode
or
IC
ACx
acoustic stimulation
A
Inferior colliculus
Spike count
p=0.05
Wt
Mbp shi/shi
B
Auditory cortex
Spike count
n.s.
Wt
Mbp shi/shi
n.s.
Wt
Mbp neo/neo
Latency increase (%)
shi/shi
neo/neo
Wt
R2=0.82
fit (poly2)
% MBP loss
Inferior colliculus
Auditory cortex
C
stim.
Mean
Rep. #
Jitter
D
E
F
Jitter (first sp std)
n.s.
G
stim.
Rep. #
Reliability
75%
H
I
J
Reliability (%)
n.s.
Wt
Mbp shi/shi
Mbp neo/neo
Auditory cortex
K
L
M
N
Firing rate (sp/s)
Time (ms)
Latency (ms)
p=0.06
Jitter (first sp std)
n.s.
Reliability (%)
Wt
Mct1+/-

**Figure S3. Basic response properties are not impaired in mice with dysmyelination or with an oligodendrocyte metabolic defect.**

Scheme illustrating position of inferior colliculus and cortex. **A-F)** Acute electrophysiology. **A)** Inferior colliculus response strength to single click stimulus was measured *in vivo*. *Mbp*$^{shi/shi}$ mice (red; n=6) showed a borderline significant increase in response strength in the IC compared to wild type littermates (black; n=8) (p=0.05). **B)** Primary cortical response strength was unchanged in *Mbp*$^{shi/shi}$ mice (left graph) (p=0.72, n=9 wild type, n=12 *Mbp*$^{shi/shi}$ animals) and in Mbp$^{neo/neo}$ mice (middle graph) (p=0.17, n=8 wild type, n=8 *Mbp*$^{neo/neo}$ animals). The latency increase for each mouse line relative to their internal wild type was correlated to the amount of *Mbp l*oss (right graph). This correlation fitted a polynomial fit of 2 degrees ($R^2$=0.82). **C)** Schematic of the jitter measurement. Jitter was defined as the standard deviation of the first spike latency/repetition. Jitter in *Mbp*$^{shi/shi}$ animals is not changed. **D)** at the IC level (p=0.065), **E)** or in the ACx (p=0.90). **F)** Similarly, *Mbp*$^{neo/neo}$ mice do not show a change in jitter (p=0.26). **G)** Schematic of the measurement of spiking reliability: the percentage of repetitions that elicited at least one spike within a 50 ms window. Reliability in *Mbp*$^{shi/shi}$ animals is not changed **H)** at the IC level (p=0.51), **I)** or in the ACx (p=0.80). **J)** *Mbp*$^{neo/neo}$ mice do not show a change in reliability in the ACx (p=0.44). All bar plots represent the mean data for all animals recorded per group and error-bars the S.E.M. In B-I) S.E.M. are represented with shaded areas. Outliers are depicted with an ‘x’ and were not considered in the statistical analysis. **K-N)** Acute electrophysiology in auditory cortex of MCT1$^{+/-}$ mice. **K)** Mean PSTH evoked by a click of 80 dB. The strength of the response was reduced in MCT1$^{+/-}$ mice (purple) compared to wild type (black). **L)** Response latency measured at the base of the PSTH is mildly but not significantly increased in MCT1$^{+/-}$ animals (p=0.06). **M)** Jitter and **N)** reliability (% repetitions that elicited at least 1 spike in a window of 37 ms) are not changed in MCT1$^{+/-}$ mice (p=0.41 and p=0.28 respectively). n=6-8 wild type mice and n=5-6 MCT1$^{+/-}$. All measurements depict the mean of the recorded animals (bar graph) and S.E.M. either with error bars or a shaded area.

### Figure S4

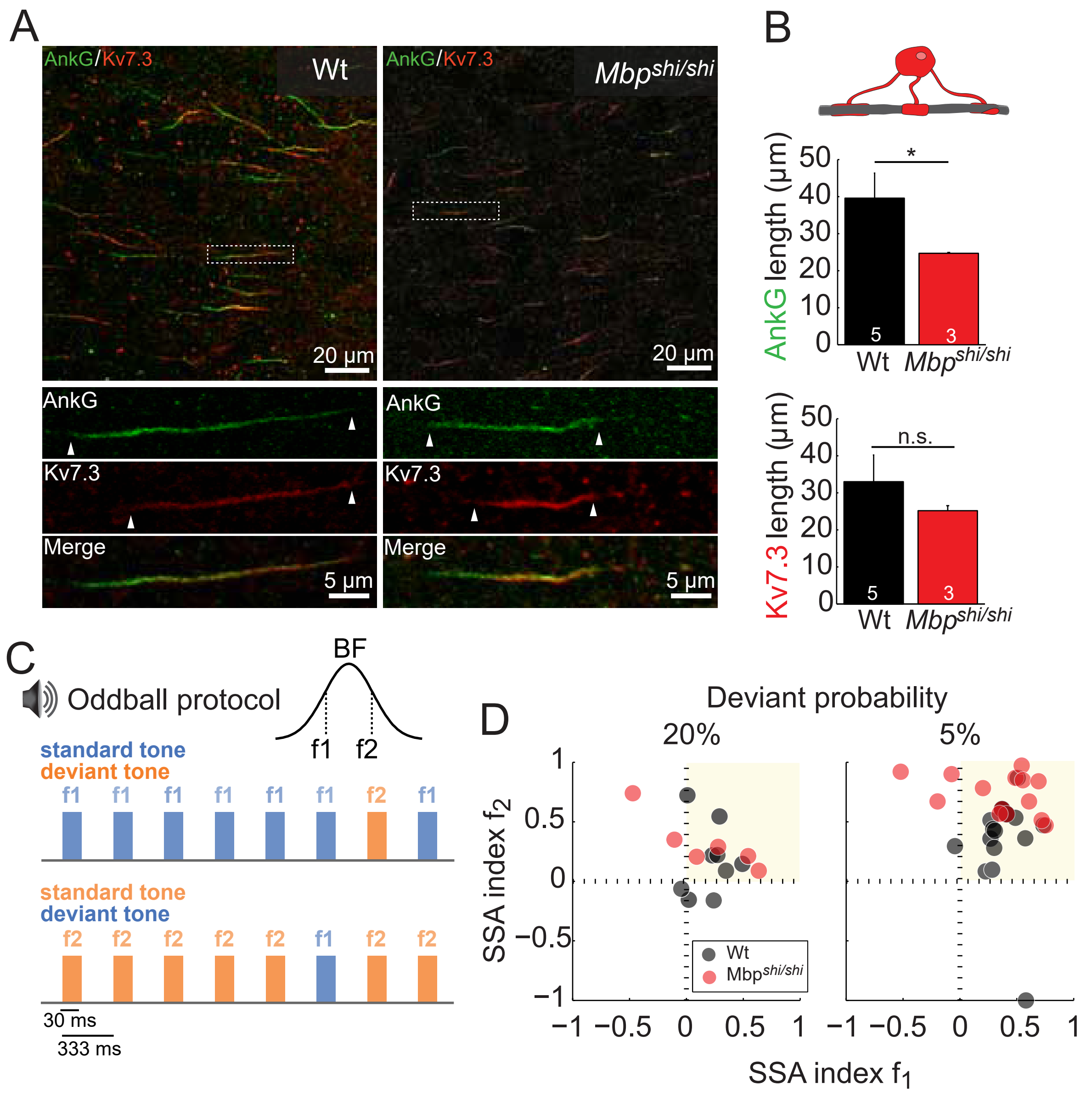

A
AnkG/Kv7.3
Wt
Mbpshi/shi
20 µm
AnkG
Kv7.3
Merge
5 µm
B
AnkG length (µm)
Kv7.3 length (µm)
*
n.s.
Wt
Mbpshi/shi
C
Oddball protocol
BF
f1 f2
standard tone
deviant tone
30 ms
333 ms
D
Deviant probability
20%
5%
SSA index f2
SSA index f1
Wt
Mbpshi/shi

**Figure S4. AIS is reduced in length with dysmyelination.**

**A)** AnkyrinG (green) and $K_v7.3$ channels (red) were stained in mouse auditory cortex (layer II/III) and used to measure AIS length in wild type (left) and *Mbp*$^{shi/shi}$ (right) mice. Below: magnifications of individual AIS stained with AnkG (green), $K_v7.3$ (red) and merge. **B)** Quantification of AnkG length revealed a significant reduction in AIS length between groups (p=0.035) while the quantification of $K_v7.3$ did not (p=0.57). For display purposes the background fluorescence, based on a ROI without an AIS, was subtracted from the whole image. n= 5 wild type mice, 20-55 AIS per mouse. n= 3 *Mbp*$^{shi/shi}$ animals, 20-30 AIS per mouse. **C)** Schematic of the oddball protocol used to test stimulus specific adaptation. Two tones differing in a Δf of 10% were presented with different probability. The 'standard' tone was presented with high probability (80 or 95%) and the 'deviant' tone with low probability (20 or 5%). Tone presentation rate was 3Hz. **D)** Stimulus specific adaptation indices (normalized difference between deviant and standard response) for the two tones were plotted against each other. Indices that fall in the upper right corner (yellow patch) reflect SSA. For low deviant probabilities (Dp=5%) indices were above 0.3, indicating that responses to a sound as deviant sounds was at least double in magnitude than to the sound as standard. This effect diminished as the probability of the deviant sound increased. While no difference in SSA was observed between the groups at Dp=20% ($p>0.35$), for Dp=5%, *Mbp*$^{shi/shi}$ mice showed even more pronounced SSA ($p<0.001$). n=9-15 Wt recordings. n=6-14 *Mbp*$^{shi/shi}$ recordings from. All plots represent the mean data of all recordings per group and error-bars the S.E.M.

### Figure S5

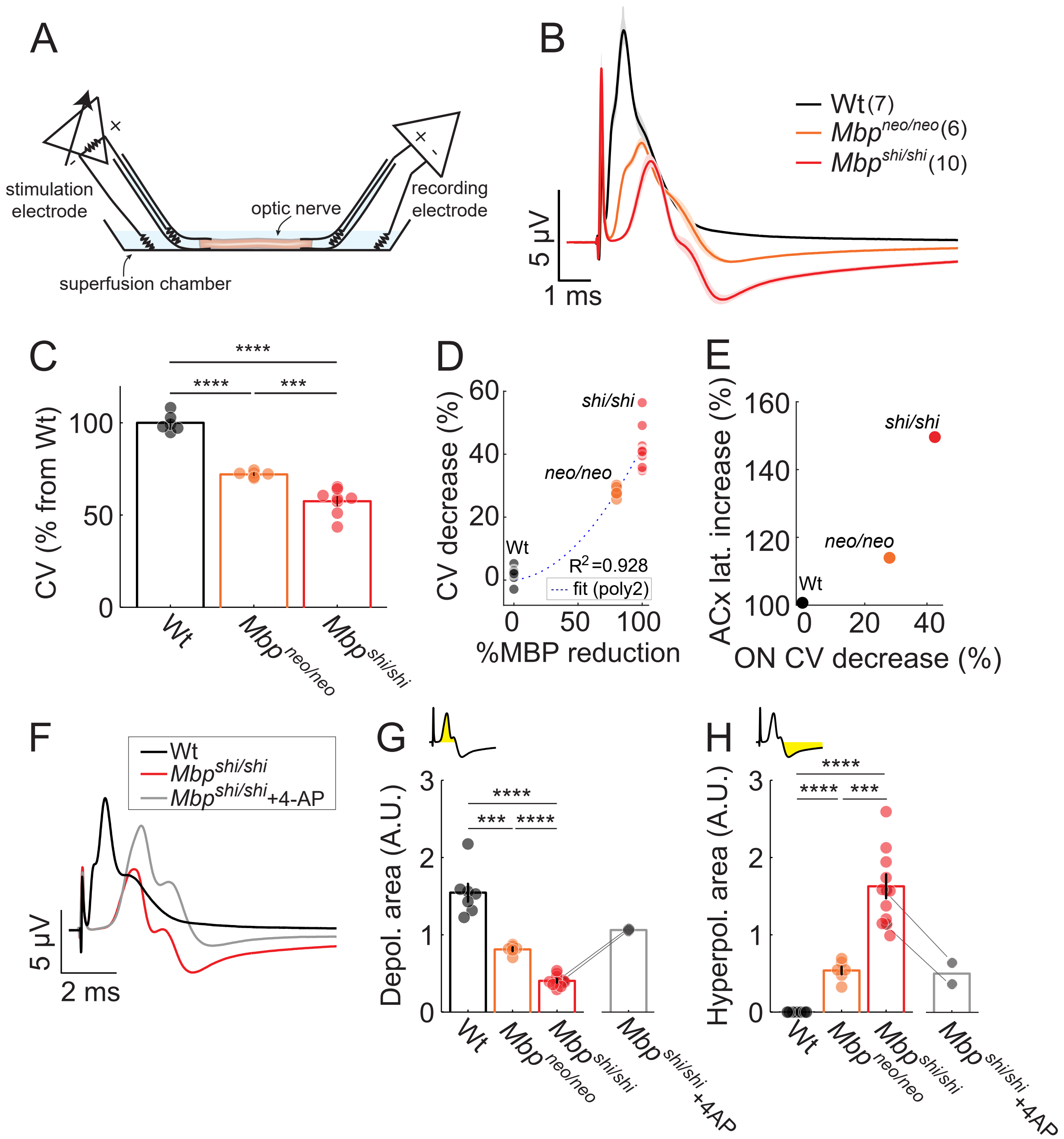

A
stimulation electrode
optic nerve
recording electrode
superfusion chamber
B
Wt(7)
Mbp neo/neo (6)
Mbp shi/shi (10)
5 µV
1 ms
C
CV (% from Wt)
Wt
Mbp neo/neo
Mbp shi/shi
D
CV decrease (%)
%MBP reduction
shi/shi
neo/neo
Wt
R2 =0.928
fit (poly2)
E
ACx lat. increase (%)
ON CV decrease (%)
shi/shi
neo/neo
Wt
F
Wt
Mbp shi/shi
Mbp shi/shi +4-AP
5 µV
2 ms
G
Depol. area (A.U.)
Wt
Mbp neo/neo
Mbp shi/shi
Mbp shi/shi +4AP
H
Hyperpol. area (A.U.)
Wt
Mbp neo/neo
Mbp shi/shi
Mbp shi/shi +4AP

**Figure S5. White matter abnormalities resulting from full and partial dysmyelination**

**A)** *Ex vivo* recordings from the optic nerve were performed in a superfusion chamber using suction electrodes. **B)** Compound action potentials (CAP) from Wt (black), *Mbp*$^{shi/shi}$ (red) and *Mbp*$^{neo/neo}$ mice (orange) are shown; stimulation current: 0.7 mA. Traces show the mean of all optic nerves; shadow around the mean shows the S.E.M. The typical triphasic CAP shape is seen in Wt animals and *Mbp*$^{neo/neo}$ mice, *Mbp*$^{shi/shi}$ show a strong reduction of the response amplitude and loss of the triphasic shape **C)** Conduction velocity measured at stimulation intensity of 0.7 mA showing a significant reduction in *Mbp*$^{shi/shi}$ nerves ($p<0.0001$), and *Mbp*$^{neo/neo}$ nerves ($p<0.0001$). the effect was significantly stronger in *Mbp*$^{shi/shi}$ ($p=0.00038$). Wt: n=6 ON from 6 mice; *Mbp*$^{neo/neo}$: n=6 ON from 3 mice; *Mbp*$^{shi/shi:}$ n= 8 ON from 7 mice. **D)** Conduction velocity decrease as a function of MBP protein reduction (see Figure S1 for *Mbp*$^{neo/neo}$) for Wt, *Mbp*$^{shi/shi}$, and *Mbp*$^{neo/neo}$ mice. A polynomial fit of 2 degrees was best ($R^2$=0.928). **E)** Response latency increase in auditory cortex was correlated to conduction velocity reduction in optic nerve. **F)** Mean CAP in the presence of 25 μM 4-AP reveals that the waveform partially recovered its shape in terms of amplitude, increase in depolarizing area and decrease in hyperpolarizing area. **G)** Depolarizing CAP area (yellow area in schematic CAP shape) was reduced in both *Mbp*$^{neo/neo}$ ($p=0.00013$) and *Mbp*$^{shi/shi}$ ($p<0.0001$). The reduction correlated with the amount of *Mbp* loss, with a significant difference between *Mbp*$^{neo/neo}$ and *Mbp*$^{shi/shi}$ ($p<0.0001$). **H)** Hyperpolarizing CAP area (yellow area in schematic CAP shape) was increased in both *Mbp*$^{neo/neo}$ ($p<0.0001$) and *Mbp*$^{shi/shi}$ ($p<0.0001$) nerves. As before the increase was proportional to the amount of myelin loss, with a significant difference between *Mbp*$^{neo/neo}$ and *Mbp*$^{shi/shi}$ ($p=0.00012$). A recovery of both the G) depolarization and H) hyperpolarization area were observed in the presence of 4-AP (gray) **F-H)** Wt: n=7 ON from 7 animals; *Mbp*$^{neo/neo}$. n=6 ON from 3 animals; *Mbp*$^{shi/shi}$*.* n=10 ON from 9 animals; 4-AP treated *Mbp*$^{shi/shi}$. n=2 ON from 2 animals. All measures show the mean and black error bars, the S.E.M.

### Figure S6

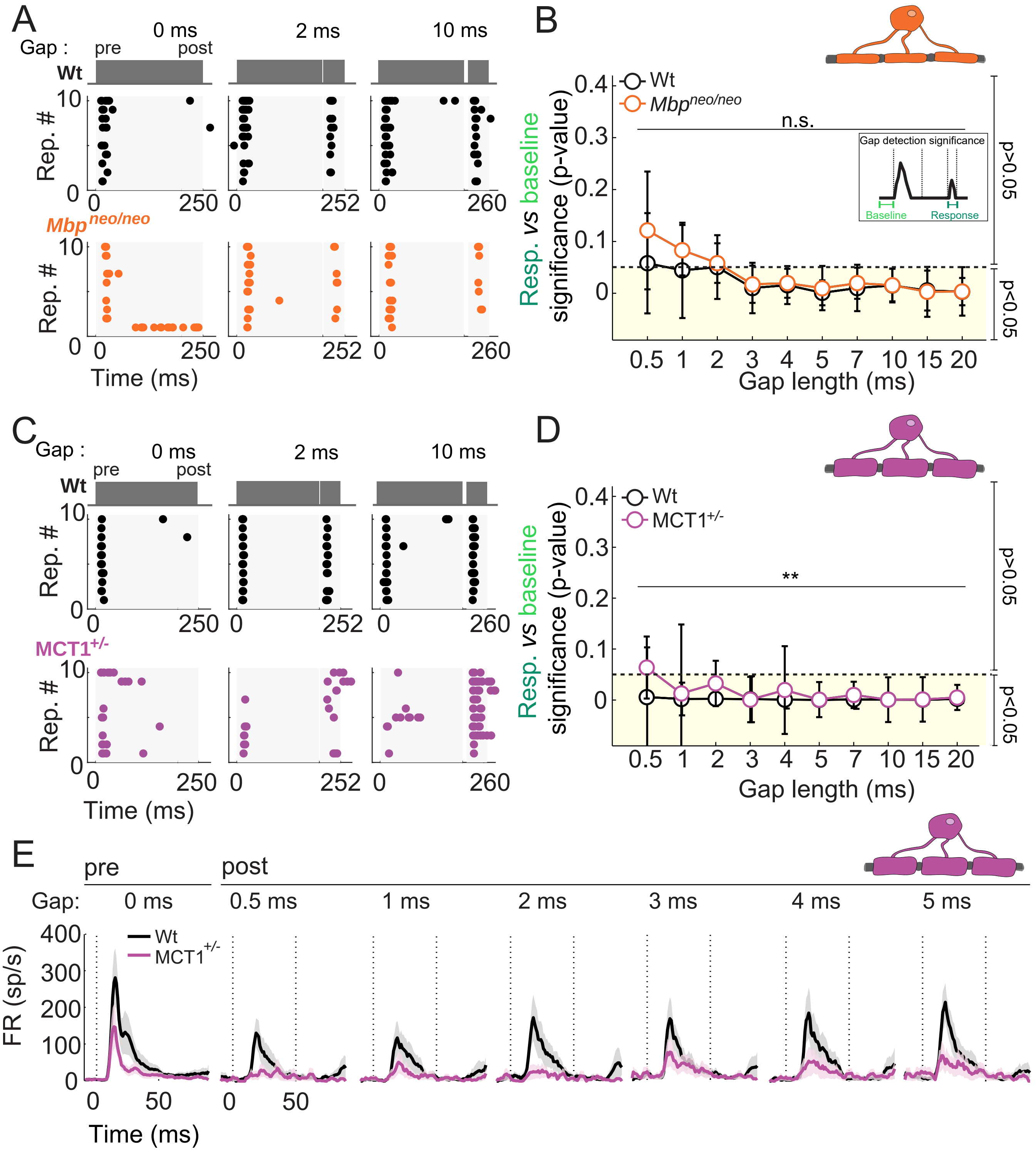
A
Gap :
0 ms
2 ms
10 ms
pre
post
Wt
Rep. #
Mbp^neo/neo
Time (ms)
B
Wt
Mbp^neo/neo
n.s.
Gap detection significance
Baseline
Response
Resp. vs baseline significance (p-value)
Gap length (ms)
p>0.05
p<0.05
C
Gap :
0 ms
2 ms
10 ms
pre
post
Wt
Rep. #
MCT1+/-
Time (ms)
D
Wt
MCT1+/-
**
Resp. vs baseline significance (p-value)
Gap length (ms)
p>0.05
p<0.05
E
pre
post
Gap:
0 ms
0.5 ms
1 ms
2 ms
3 ms
4 ms
5 ms
Wt
MCT1+/-
FR (sp/s)
Time (ms)

**Figure S6. Gap detection *Mbp*$^{neo/neo}$ and MCT1$^{+/-}$**

**A)** Individual raster plot examples of ACx spikes (dots) evoked during sound presentation (gray patches) before (pre) and after (post) silent gaps of 0 ms (left), 2 ms (center), and 10 ms (right) length in a Wt (upper, black) and a *Mbp*$^{neo/neo}$ (lower, orange) mouse during 10 stimulus repetitions (y axis) per gap length. **B)** Quantification of gap-detection significance (as in Figure 5C). The statistical difference (p-value) between baseline (50 ms window before pre-BBN) and post-gap activity per recording was obtained and the median per gap plotted. Values below 0.05 indicate successful detection. No significant effect of gap was observed between Wt (black; n=17-22 recording sites of 7 animals) and *Mbp*$^{neo/neo}$ (orange; n=13-17 recordings of 8 mice) (p=0.37). **C)** Same as A) for MCT1$^{+/-}$ mice. **D)** Same as B) for MCT1$^{+/-}$ mice. Gap-detection was successful across gap lengths for MCT1$^{+/-}$ mice (purple; n=8-9 recordings from 6 MCT1+/- mice), and Wt mice (black; n=13 recording sites from 8 animals) despite a significant difference between the groups (p=0.0027). In B) and D) dots show the median per group per gap length, and error bars the standard error of the median. Dotted line: threshold at 0.05. Yellow shadow: significant gap-detection. **E)** Average PSTH across animals for Wt (black; n=8) and MCT1$^{+/-}$ (purple; n=6) with S.E.M. as shaded area. Left to right: pre-gap responses followed by post-gap responses for 0.5 to 5 ms gaps. Dotted vertical lines at time 0 ms depict the start of the sound, and at time 50 ms the end of the post BBN.

### Figure S7

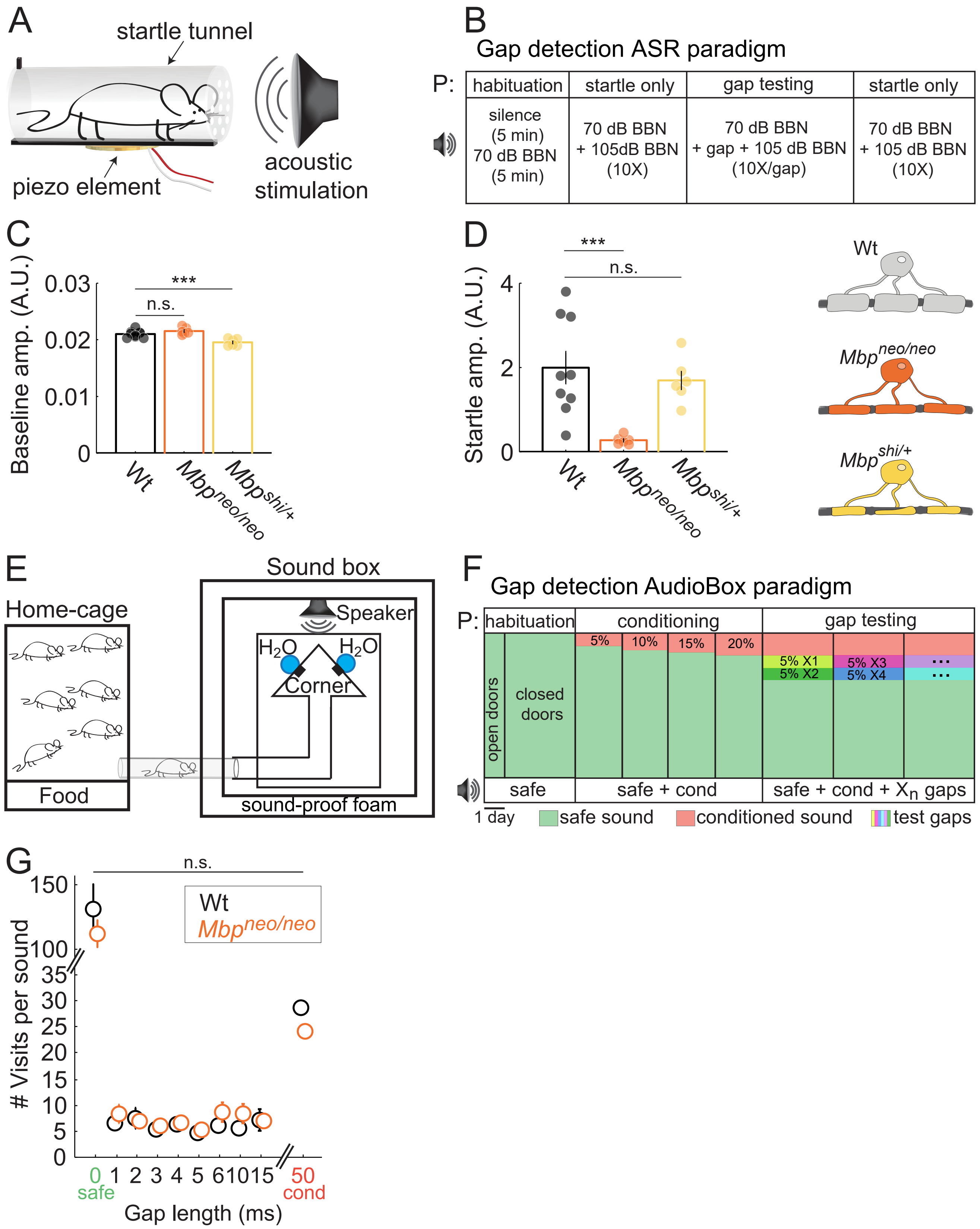
A
startle tunnel
piezo element
acoustic stimulation
B
Gap detection ASR paradigm
P:
habituation
startle only
gap testing
startle only
silence (5 min) 70 dB BBN (5 min)
70 dB BBN + 105dB BBN (10X)
70 dB BBN + gap + 105 dB BBN (10X/gap)
70 dB BBN + 105 dB BBN (10X)
C
Baseline amp. (A.U.)
0.03
0.02
0.01
0
n.s.
***
Wt
Mbpneo/neo
Mbpshi/+
D
Startle amp. (A.U.)
4
2
0
***
n.s.
Wt
Mbpneo/neo
Mbpshi/+
Wt
Mbpneo/neo
Mbpshi/+
E
Home-cage
Food
Sound box
Speaker
H2O
H2O
Corner
sound-proof foam
F
Gap detection AudioBox paradigm
P:
habituation
conditioning
gap testing
open doors
closed doors
5%
10%
15%
20%
5% X1
5% X2
5% X3
5% X4
safe
safe + cond
safe + cond + Xn gaps
1 day
safe sound
conditioned sound
test gaps
G
n.s.
### Visits per sound
150
100
35
30
25
20
15
10
5
0
Wt
Mbpneo/neo
0 safe
1 2 3 4 5 6 10 15
50 cond
Gap length (ms)

**Figure S7. Behavioral temporal acuity testing protocols**

**A)** Schematic of gap-detection inhibition of the auditory startle reflex (GDIASR) test. Briefly, a mouse is introduced in the startle tunnel. A piezoelectric element below measures movement including startle elicited by loud sounds. **B)** A diagram of the GDIASR sound protocol. The experiment is divided in 4 phases. During habituation, the mouse was acclimated in the tunnel for 5 min. in silence and for a further 5 min. with a 70 dB borad-band noise (BBN) as background sound. During the 'startle only' phase, 10 startle sounds (105 dB BBN, 20 ms long) were played interspersed in the background sound to assess the pure startle response. In the 'gap testing' phase, small silent gaps of varying lengths are embedded in the background sound 50ms before the startle sound. Finally, another startle-only phase with 10 startle sounds is used to assess the level of auditory habituation. **C)** Baseline movement was not different for Wt (black, n=9) and *Mbp*$^{neo/neo}$ mice (orange; n=6) (p=0.14). A reduction in the baseline movement of *Mbp*$^{shi/+}$ (yellow; n=6) was observed (p=0.00039). **D)** Amplitude of the startle-only response (5 largest trials of the initial 10 ASR-only trials) was significantly lower in *Mbp*$^{neo/neo}$ compared to Wt (p=0.00079), but not different between Wt and *Mbp*$^{shi/+}$ (p=0.86). **E)** Schematic diagram of the AudioBox paradigm. Briefly, food is available in a 'home-cage' and water, in the 'sound box', where a corner contains water bottles. Sounds are played each time a mouse enters the corner, and access to water depends on the sound played. **F)** Diagram of the AudioBox gap-detection paradigm. During the habituation phase, mice hear the 'safe sound', a continuous 70 dB BBN, every time they enter the corner. Nose-poking in the presence of this sound triggers door opening and access to water. During the conditioning phase the conditioned sound (BBN + 50 ms silent gap) is introduced in progressively larger proportion of visits to the corner (5% to 20%). During the gap testing phase, every 2-3 days 2 new visit types are introduced, each with a different gap length, to assess gap-detection. **G)** Mean visits per sound for the duration of the experiment for Wt (black) and *Mbp*$^{neo/neo}$ (orange) animals. No differences between groups were seen for the overall sounds analyzed (p=0.29).
